# An atlas of transcriptional dynamics in maternal blood over the course of healthy pregnancy

**DOI:** 10.64898/2026.03.30.715300

**Authors:** Bjarke Feenstra, Freja Randrup Dahl Hede, Brian D. Piening, Line Skotte, Katerina Nastou, Liang Liang, João P. S. Fadista, Marie-Louise H. Rasmussen, Nikolai Madrid Scheller, Chao Jiang, Francesco Vallania, Eric Wei, Qing Liu, Hassan Chaib, Frank Geller, Heather A. Boyd, Michael P. Snyder, Mads Melbye

**Author notes:** Equal contribution (Bjarke Feenstra, Freja Randrup Dahl Hede, Brian D. Piening). These authors jointly supervised the work (Heather A. Boyd, Michael P. Snyder, Mads Melbye).

## Abstract

Pregnancy results in profound physiological changes driven by dynamic and precisely programmed molecular processes. Maternal peripheral blood is generally the specimen of choice for studying these processes, as it is easily accessible and essential for many aspects of maintaining a healthy pregnancy. Here, we present a high-resolution atlas of the dynamic temporal changes in the transcriptome of maternal peripheral blood in healthy human pregnancy. We generated comprehensive RNA sequencing data in 802 weekly samples from 31 healthy pregnant women from the first trimester until after delivery. Using a strict discovery and replication setup, our longitudinal analysis of gene expression identified 720 genes with robust pregnancy-specific expression patterns. Using weighted graph correlation network analysis, we identified nine pregnancy-associated transcriptional modules that reveal a strong, coordinated enrichment of innate/neutrophil and antiviral immune programs, alongside changes in adaptive immunity (T cell differentiation and signaling), erythropoiesis and hemoglobin metabolism. Cell-type deconvolution revealed that these transcriptomic shifts were accompanied by increased relative neutrophil proportions and reduced naive CD4 and CD8 T cells in pregnancy. We provide a comprehensive characterization of dynamic changes across pregnancy, highlighting maternal blood as a key systemic regulator in healthy gestation. Together, our findings establish a reference atlas of healthy pregnancy, which can be used to identify dysregulated processes and mechanisms in women with pregnancy complications.

**Graphical abstract:** 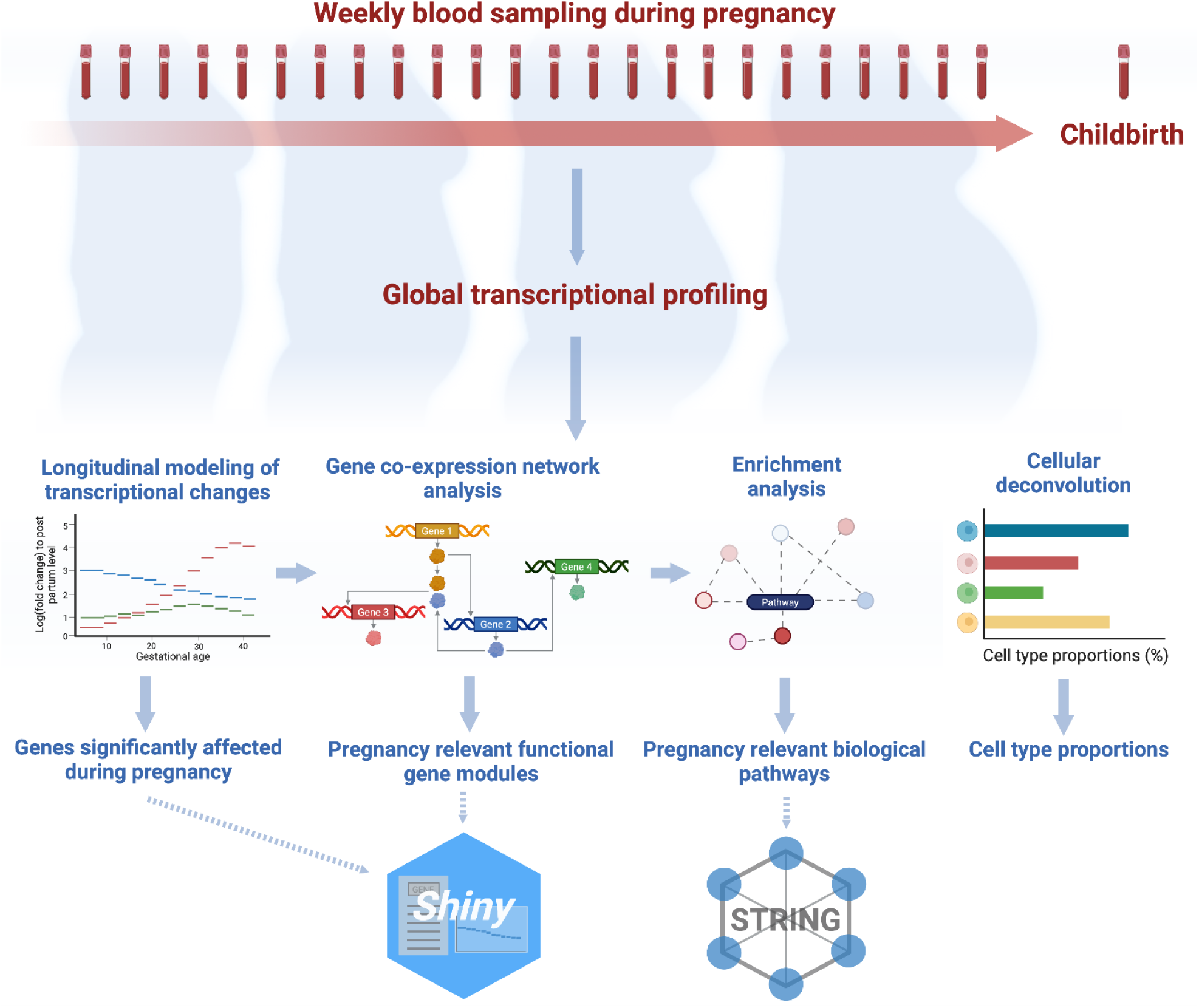

- 720 genes showed robust pregnancy specific expression patterns.
- Co-expression analysis clustered the genes into nine modules with distinct dynamics.
- Enrichment in pathways involved in innate and neutrophil-mediated immunity, antiviral responses, T cell differentiation and signaling, erythropoiesis and hemoglobin metabolism.
- Cell-type deconvolution showed increases in neutrophils and decreases in naïve CD4 and CD8 T cells.
- The atlas of detailed longitudinal transcriptional changes provides a baseline reference for healthy pregnancy.
- Results for all genes and protein-protein interaction networks are made available for interactive exploration.

## Introduction

Healthy pregnancy is characterized by tightly regulated, ordered physiological processes that ensure normal fetal development and delivery after a well-defined period of gestation. Dysregulation of these processes can produce adverse consequences for the health of both mother and fetus. Much effort has been devoted to identifying molecular processes disrupted in women with pregnancy complications such as preeclampsia, placental disorders, fetal growth restriction, hyperemesis, and preterm delivery^1–5^, whereas ‘omics research into the healthy pregnancy has received less attention. To fill this gap, we characterized the dynamic transcriptional changes in maternal peripheral blood with high temporal resolution over the course of healthy human pregnancy.

Sampling of maternal peripheral blood is a minimally-invasive and routinely-performed procedure with little risk to either the pregnant woman or the fetus, making it highly feasible and ethically appropriate for large-scale studies and clinical testing. Maternal blood provides a readily available source of circulating biomarkers, leukocytes and cell-free DNA and RNA, which together capture systemic gene expression signatures. High-throughput transcriptomic profiling in maternal blood using RNA sequencing therefore enables comprehensive analyses of gene expression dynamics across gestation.

Few previous studies have measured gene expression in the maternal circulation at multiple time points during normal pregnancy. A microarray-based gene expression study of 49 women included samples at up to six time points^6^ and an RNA-sequencing study of 14 healthy women included samples at three time points in pregnancy along with pre- and post-pregnancy samples^7^. These previous studies highlighted specific genes with pregnancy-associated changes in transcription, and patterns of change in immune-related gene sets and in cell-type proportions.

In this study, we leverage dense, repeated sampling of maternal blood across gestation to build a high temporal resolution longitudinal transcriptomic resource. This design enables within-individual dynamic modeling of gene expression trajectories, capturing temporal regulatory patterns that are difficult to resolve in cross-sectional cohorts. To our knowledge, this dataset represents the first cohort with weekly whole-transcriptome profiling throughout pregnancy, providing the resolution necessary to detect subtle and transient transcriptional changes occurring during healthy pregnancy. These data establish a reference for healthy pregnancy, providing both a biologically informative foundation for exploring and understanding the underlying mechanisms of gestation and a comparative framework for integrating complementary data types, including those generated from complicated pregnancies.

## Results

### RNA-sequencing of weekly maternal blood samples in pregnancy

We characterized the dynamic biological processes of pregnancy with high temporal resolution in a cohort of healthy pregnant Danish women who had blood drawn weekly from the first trimester until after delivery. We performed RNA-sequencing of cellular RNA from 802 blood samples from 31 participants, divided into a discovery set (531 samples, 21 women) and a replication set (271 samples, 10 women), corresponding to almost 26 time points per woman on average (**Fig. 1**). A flow chart of the study design is shown in **Supplementary Fig. 1.**

**Fig. 1:**
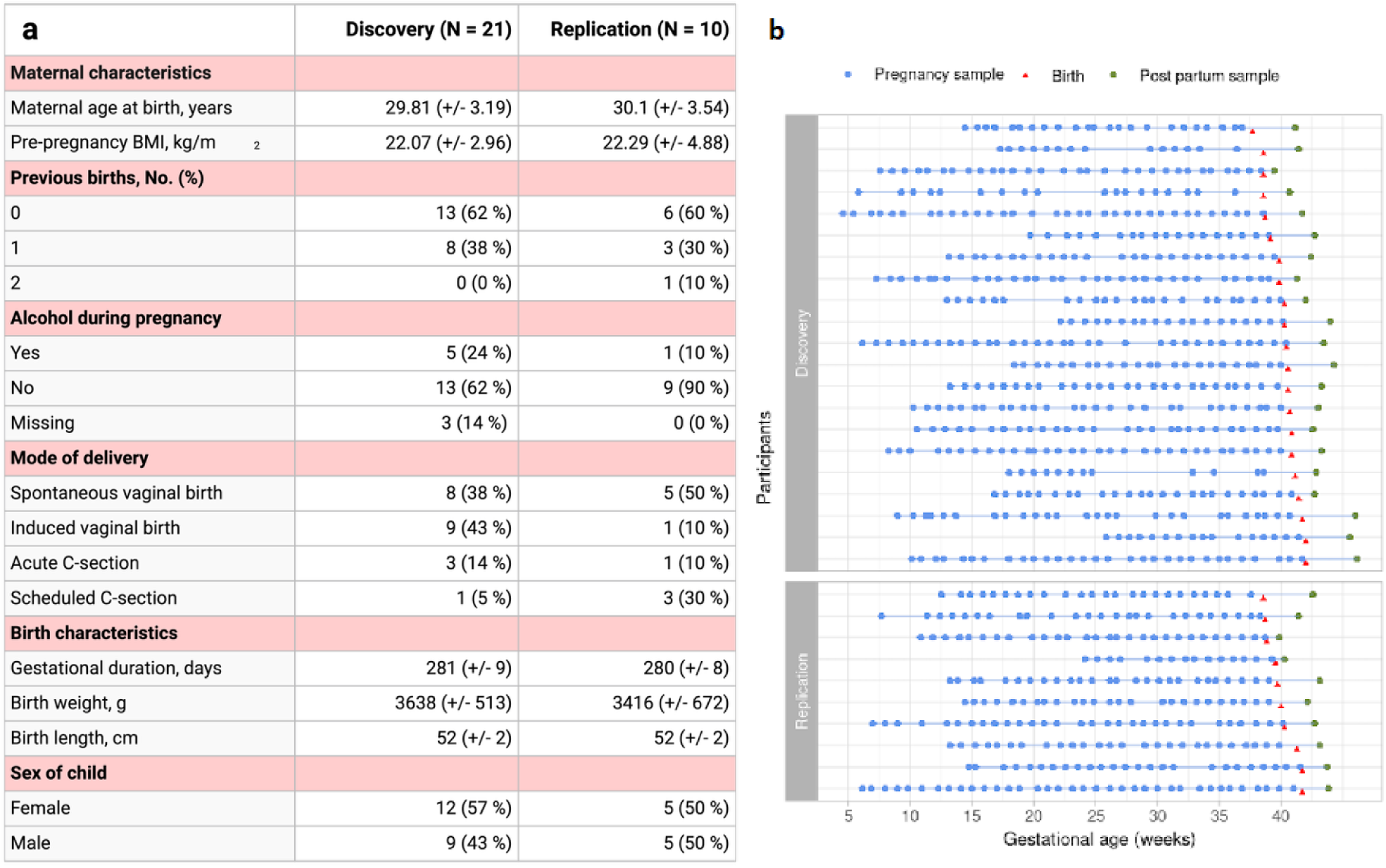
Characteristics of the study sample. **a,** Table of characteristics of the discovery and replication sets. Values are presented as means (+/- standard deviation) or counts (%) where applicable. No participants were smoking during their pregnancy. **b,** Gestational week at sample collection for each group during pregnancy (blue circles) and postpartum (green circles). Gestational age at delivery is also shown (red arrows).

RNA-sequencing of the discovery samples averaged 29.1 million reads (71.3% uniquely mapped), while replication samples averaged 20.6 million reads (50.9% uniquely mapped). After removing 551 rRNA-annotated transcripts, retaining repeatedly detected transcripts, and restricting to uniquely assigned reads, we retained 32,104 transcripts in discovery and 24,967 in replication for downstream analyses (See **Supplementary Text** for details).

### Longitudinal modelling of expression levels

The dense sampling scheme allowed for detailed longitudinal modeling of the transcriptome across pregnancy. Using a longitudinal log-linear mixed effects model accounting for the random effects resulting from differences between the study participants, we identified 4255 transcripts in the discovery cohort with expression levels varying significantly (*P* < 1.6 x 10^-6^, i.e. Bonferroni corrected for 32,104 transcripts in the discovery batch) through pregnancy. Additionally, the median *P* value in the discovery cohort was 0.037, indicating that the relative expression levels of most transcripts are affected during pregnancy.

Among the 4,255 transcripts that were significantly associated in the discovery cohort, pregnancy-associated patterns of expression were robustly validated in the replication cohort and in the combined cohorts for 720 transcripts (**Supplementary Table 1)**. This resulted in a conservative set of transcripts with consistent variation in transcription levels across the pregnant women compared to postpartum levels (referred to as “pregnancy-associated” from here). This set contained genes showing patterns of both gradually increased expression (e.g., *OLFM4*, combined *P = 9.39 × 10^−146^*), decreased expression (e.g., *ABCG1*, combined *P = 1.84 × 10^−140^*) and relatively stable levels of increased expression (e.g., *ANXA3*, combined *P = 2.07 × 10^−109^*) throughout gestation compared to the postpartum level (**Fig. 2**). To facilitate further exploration, we provide results for all analyzed transcripts in an interactive interface (See **Data Availability**).

**Fig. 2:**
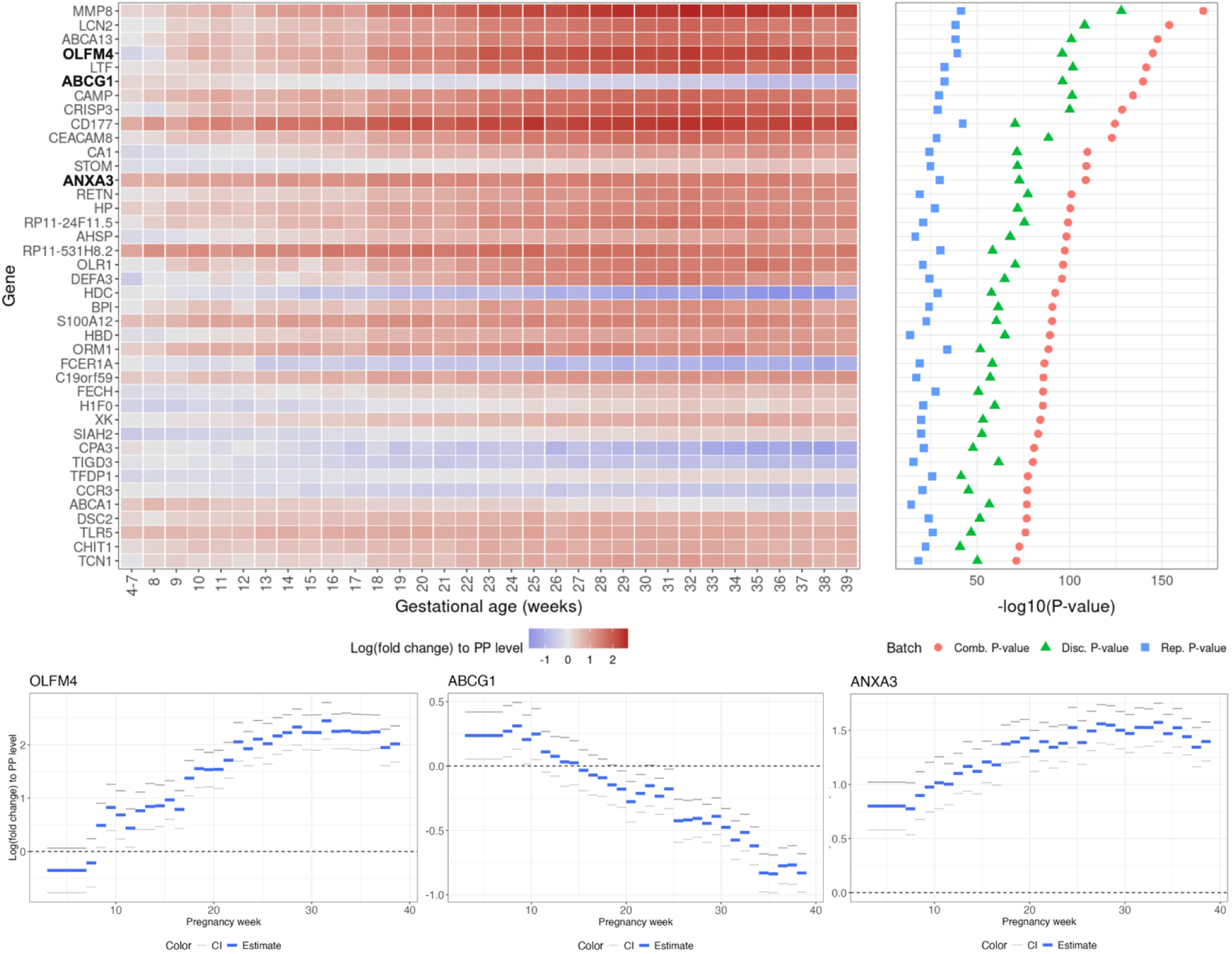
The 40 genes most significantly associated with pregnancy. **a,** Heatmap showing estimates of log fold changes of gene expression levels compared to the postpartum (PP in figure) reference level by pregnancy week. Shown are the 40 transcripts with the lowest combined *P* values among the 720 robustly replicated transcripts. **b,** Corresponding -log10 *P* values from the discovery (green), replication (blue), and combined (red) samples. **c,** Examples of genes with representative trajectories: gradually increasing expression (*OLFM4*), decreasing expression (*ABCG1*), and relatively stable increased expression (*ANXA3*) during gestation compared to the postpartum reference. These genes are highlighted in bold in the heatmap.

Among the 720 replicated transcripts, 340 were previously reported to be associated with pregnancy in studies with much sparser sampling, based on RNA-sequencing^7^ or microarray analysis^6^ (**Supplementary Fig. 2**, **Supplementary Table 2**).

### Enrichment of immune-related processes, pathways, functions and cell components

The STRING database^8^ was used to construct functional association networks and to perform functional enrichment analysis of the 720 pregnancy-associated transcripts (which mapped to 664 Ensembl^9^ Protein IDs, see **Supplementary Table 3**). STRING uses one protein per gene, which is why they are used interchangeably below^8^. Gene Ontology (GO)^10^ enrichment analyses revealed overrepresentation of terms related to immune- and stimulus-response processes, molecular functions related to binding and activity of, e.g., immune receptors, and metabolic enzymes, and various cellular components such as vesicles and granules. Reactome^11^ and Kyoto Encyclopedia of Genes and Genomes (KEGG)^12^ analyses further highlighted pathways related to immune activation (**Extended Data Fig. 1, Extended Data Table 1**).

### Pregnancy-associated transcripts cluster into nine modules based on co-expression patterns

Next, we used weighted graph co-expression network analysis (WGCNA)^13^ to resolve the 720 replicated transcripts into nine modules (**Supplementary Table 1**). Each module represents a set of transcripts with correlated expression patterns throughout pregnancy across samples, except for the grey module, which contained 46 transcripts that did not group with others in terms of co-expression. We investigated the log fold change estimates for each of the co-expression modules, which displayed distinctive patterns of dynamic change during pregnancy (**Fig. 3**).

**Fig. 3:**
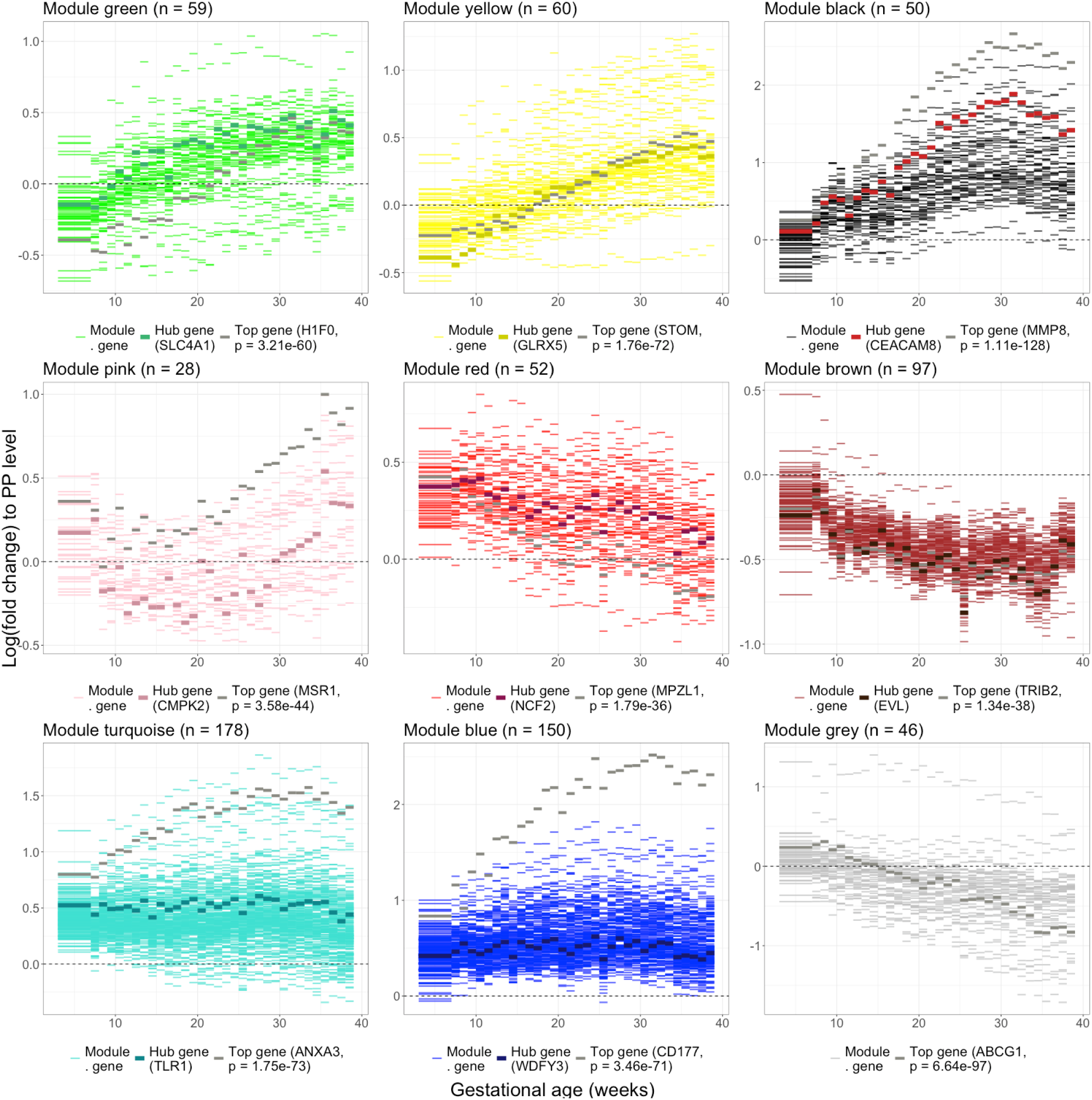
Expression dynamics of nine gene co-expression modules. Each panel displays the log fold change in expression in samples taken weekly throughout pregnancy compared to the postpartum expression levels for each module. Transcripts with the smallest discovery *P* values are highlighted in darker gray, while hub transcripts (highest module connectivity) are shown in a darker shade of the module color (dark brown for the black module).

We then performed enrichment analysis using the STRING database to biologically annotate each co-expression module (**Fig. 4**, for STRING Permalinks see **Extended Data Table 1**). Results for the individual modules are described in the following subsections.

**Fig. 4:**
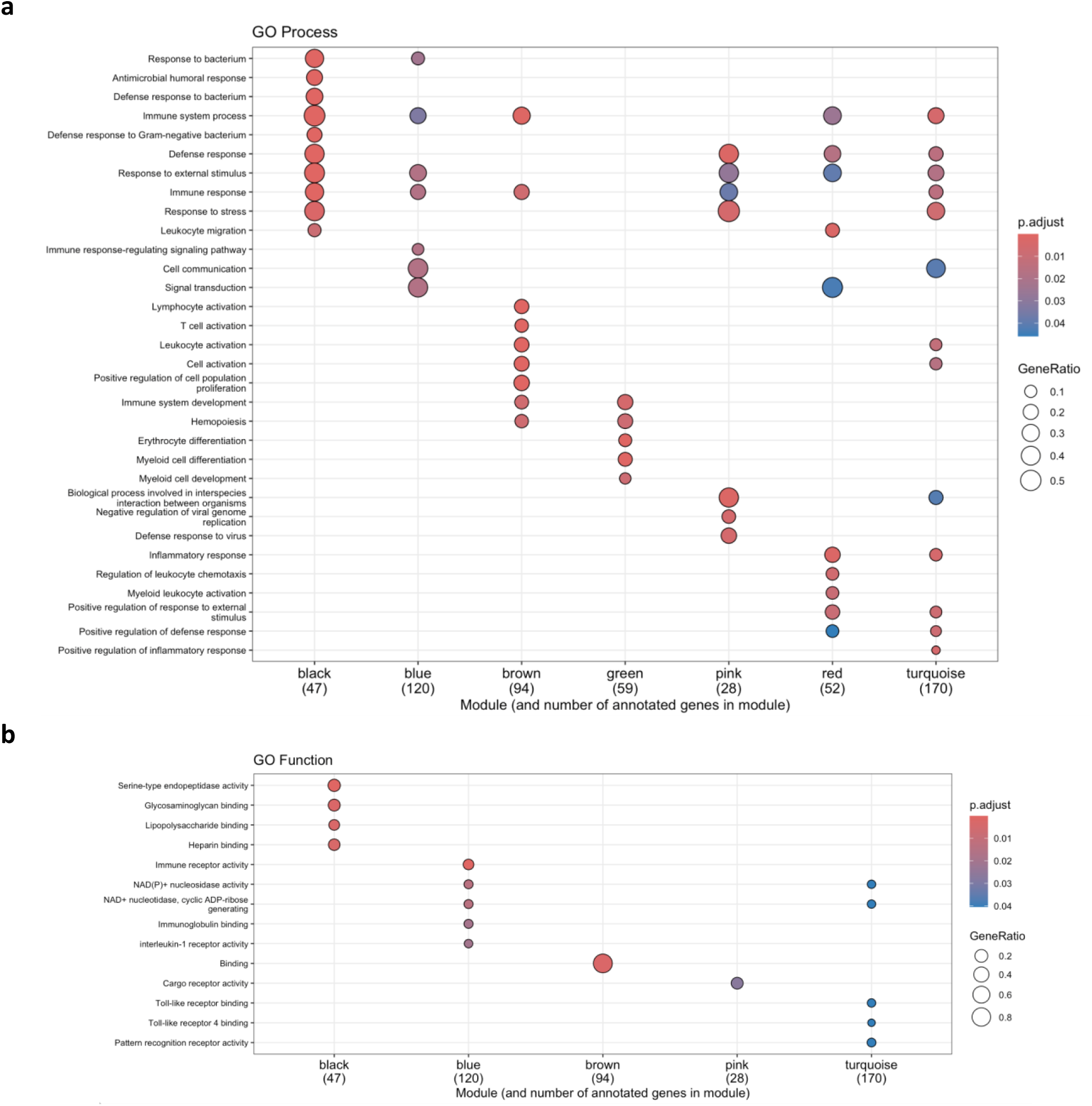
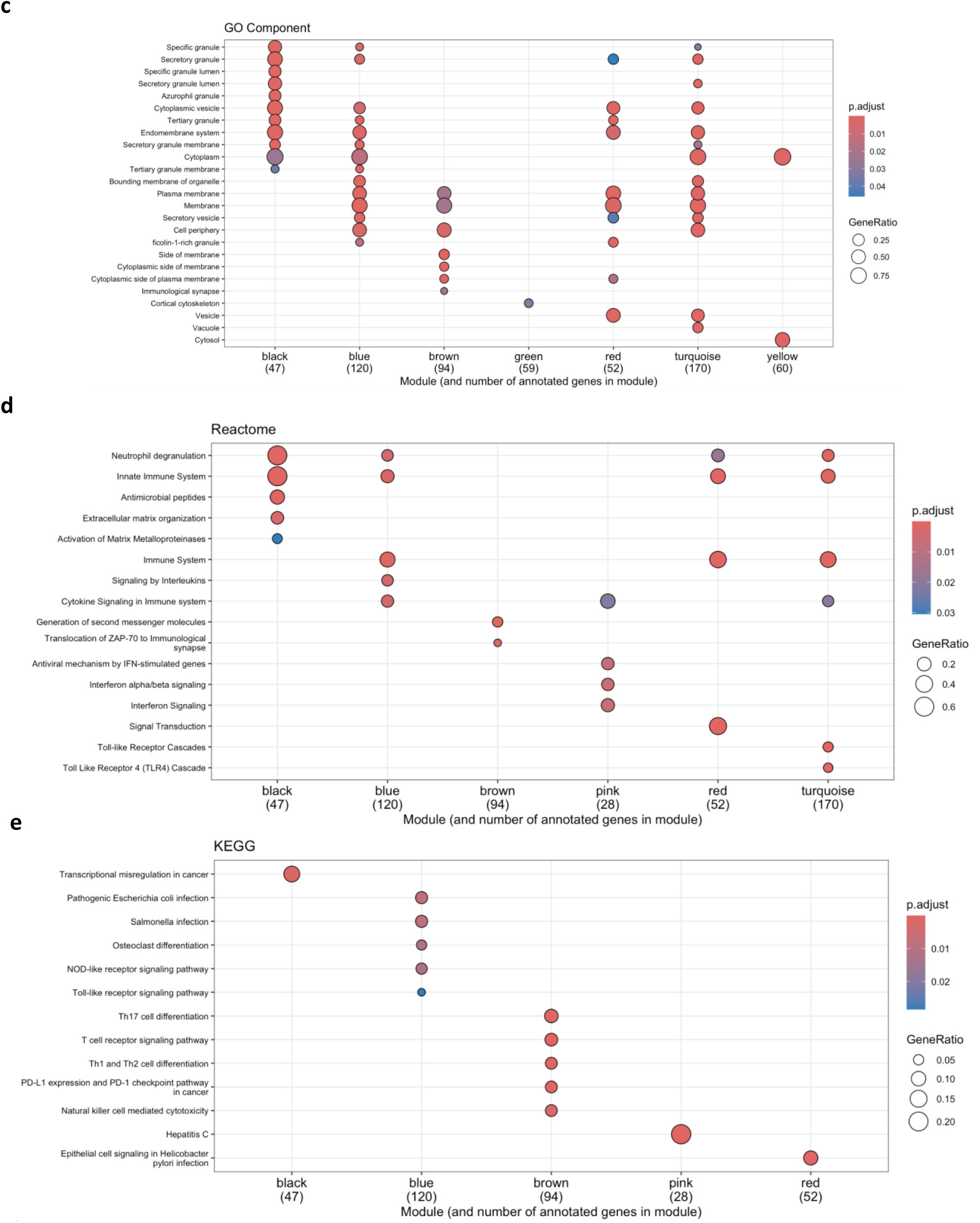
Enrichment analyses for the nine co-expression modules. Dot plots illustrate the magnitude and significance of the top enriched terms obtained via the STRING database for the nine co-expression modules on the x axis. The panels show enrichment terms related to **a,** GO Biological Process, **b,** GO Molecular Function, **c,** GO Cellular Component, **d,** Reactome Pathways, and **e,** KEGG pathways. The module labels on the x axis indicate the number of annotated proteins from each module included in the enrichment analysis. Modules with no significantly enriched terms in a given category are omitted from the relevant panel.

### Modules showing patterns of gradual increase in expression: cell proliferation and differentiation, innate immunity, and antiviral response

Four modules comprising 197 transcribed genes showed gradual increases during pregnancy relative to the postpartum reference levels but differed in their temporal profiles and biological characteristics.

The green module (**Fig. 3, Extended Data Fig. 2**) was enriched for terms related to cell proliferation and differentiation, and localized to the cortical cytoskeleton (**Fig. 4**), suggesting involvement in structural or developmental processes. The top-scoring functional associations in the green module were related to genes involved in erythropoiesis and heme biosynthesis, including *FECH*, *EPB42*, *ALAS2*, *SLC4A1*, *ANK1*, *TRIM58*, and *SPTB* (**Extended Data Fig. 2**).

In the yellow module, expression levels were initially lower than postpartum levels followed by a progressive increase during pregnancy (**Fig. 3, Extended Data Fig. 3**). The module was enriched primarily for terms related to the cytoplasmic compartment (**Fig. 4**). Few high-confidence functional associations were seen in the yellow module. Observed associations involved the genes *CA1*, *CA2*, *HBP*, and *AHSP*, which are implicated in CO₂ transport and pH regulation in erythrocytes, and hemoglobin formation (**Extended Data Fig. 3**).

The black and pink modules displayed clear immune-related signatures. The black module (**Fig. 3**, **Fig. 5**) was linked to innate immunity, with enrichment in processes such as response to bacterial and external stimuli and neutrophil degranulation, as well as localization to vesicles and granules, organelles typical of neutrophils and other innate immune cells^14^. Enriched terms included serine-type endopeptidase activity and binding of glycosaminoglycans and lipopolysaccharides (**Fig. 4**). The transcripts with the lowest *P* values were predominantly found in the black module and included *MMP8*, a neutrophil-derived matrix metalloproteinase involved in extracellular matrix remodeling and inflammatory processes, including cervical ripening^15^. The black module showed highly interconnected protein associations including *MMP8*, *LCN2*, *LTF*, *CAMP*, *CEACAM8*, *ELANE*, *DEFA3*, *DEFA4*, *PRTN3*, *AZU1* and many other genes with a range of roles related to neutrophil degranulation and inflammatory signaling (**Fig. 5**). The expression dynamics of the black module showed increasing gene expression until approximately 5-10 weeks prior to parturition, followed by a small drop in expression levels.

**Fig. 5:**
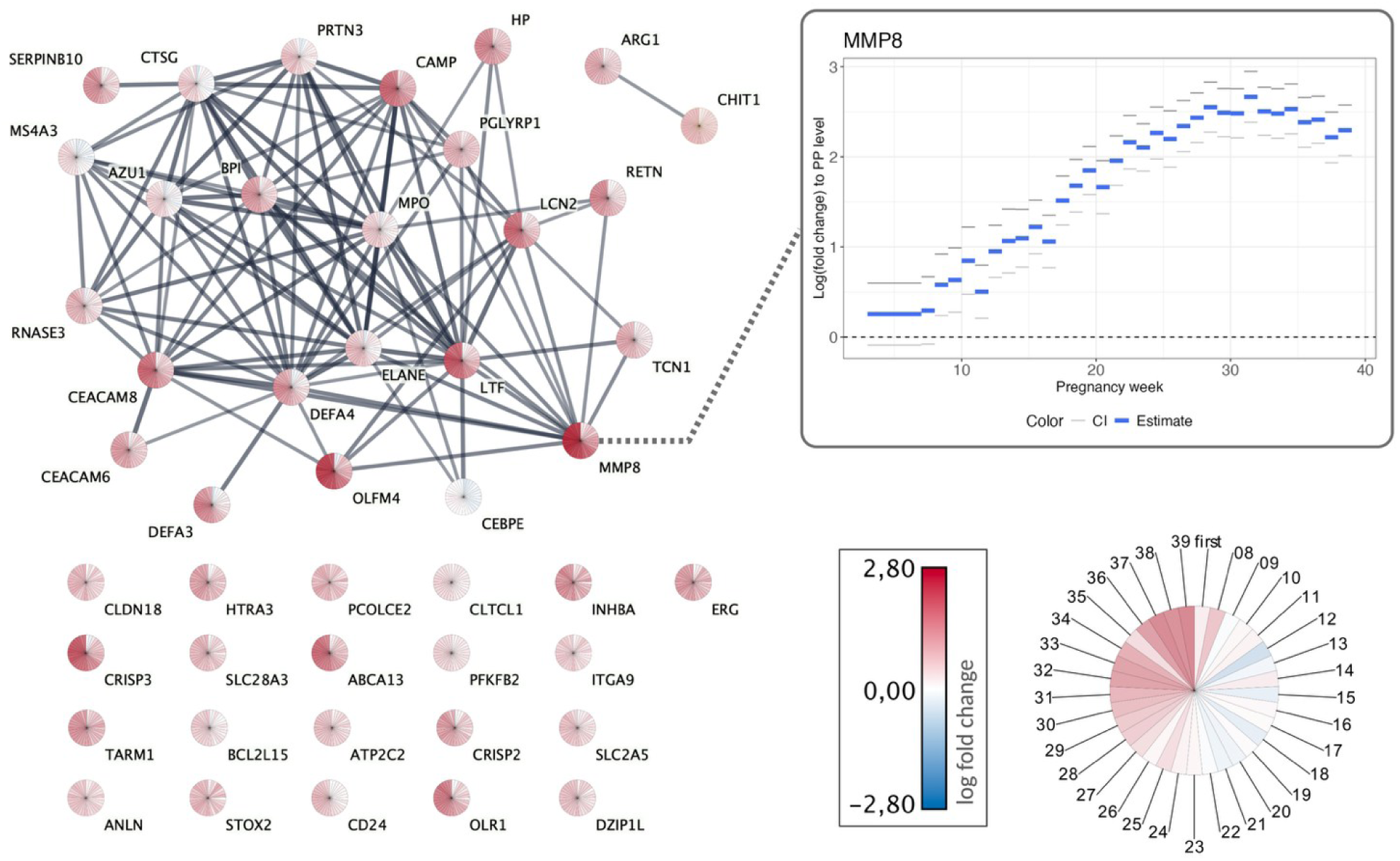
Functional associations among genes co-expressed in the black module. Nodes represent individual genes, and edges reflect STRING database interactions with high confidence (score ≥ 0.7). Each node is visualized as a pie chart, where each segment corresponds to a pregnancy week from gestational weeks 4-7 (first), then to 8 and up to 39. The shading within each segment reflects the log fold change estimates in the corresponding week, mapped using a continuous blue to red color gradient for low to high expression. This radial representation highlights the temporal dynamics of gene expression across pregnancy at single-gene resolution. The highlighted node represents *MMP8*, the module’s top gene in the discovery analysis. The plot shows its log fold change during pregnancy compared to the postpartum level with 95% confidence intervals.

In contrast, expression levels of genes in the pink module (**Fig. 3, Extended Data Fig. 4**) initially decreased during the first 10-15 weeks of pregnancy, followed by a sustained increase. This module was enriched for antiviral and interferon responses, with associated terms including response to viral and external stimuli, immune signaling, and cargo receptor activity, indicating a role in host-pathogen interactions and intracellular trafficking (**Fig. 4**). High-confidence functional associations were annotated among *IFI44L*, *IFI6*, *IFIT1*, *MX1*, *OAS3*, *EIF2AK2*, *CMPK2*, *PARP12*, and *SIGLEC1*, comprising interferon-stimulated genes that are involved in type I interferon-mediated antiviral response (**Extended Data Fig. 4**).

### Modules showing patterns of gradually decreased expression: innate and T cell immunity

The red and brown modules exhibited overall decreasing expression levels as pregnancy progressed, although there were brief increases in expression in the brown module a few weeks prior to parturition (**Fig. 3**). Expression levels in the red module were generally higher and the brown module generally lower than the postpartum reference levels (**Fig. 3**). Both modules showed associations with immune processes.

The red module (**Extended Data Fig. 5**) was enriched for terms related to innate immunity (**Fig. 4**), including leukocyte activation, inflammatory response, and leukocyte migration, as well as neutrophil-mediated functions, such as neutrophil degranulation and localization to membrane, vesicles, and ficolin-1-rich granules, which are present in human neutrophils^16^. Functional associations in the red module involved the tyrosine-protein kinases *LYN* and *SYK* which play roles in B cell response regulation, and *FPR1*, *FPR2*, and *CXCR2*, which encode receptors for neutrophil chemotactic factors (**Extended Data Fig. 5**).

The brown module (**Fig. 6**) was enriched for terms related to T cell activity, including lymphocyte activation and cell proliferation, as well as several T cell-related KEGG pathways such as differentiation of T-helper (Th) cells of type Th1, Th2, and Th17, and the T cell receptor signaling pathway. The top-scoring functional associations in the brown module were between *LCK* and other genes with roles in regulating of T cell receptor activity, including *GRAP2*, *ZAP70*, *CD3E* and *CD247* (**Fig. 6**).

**Fig. 6:**
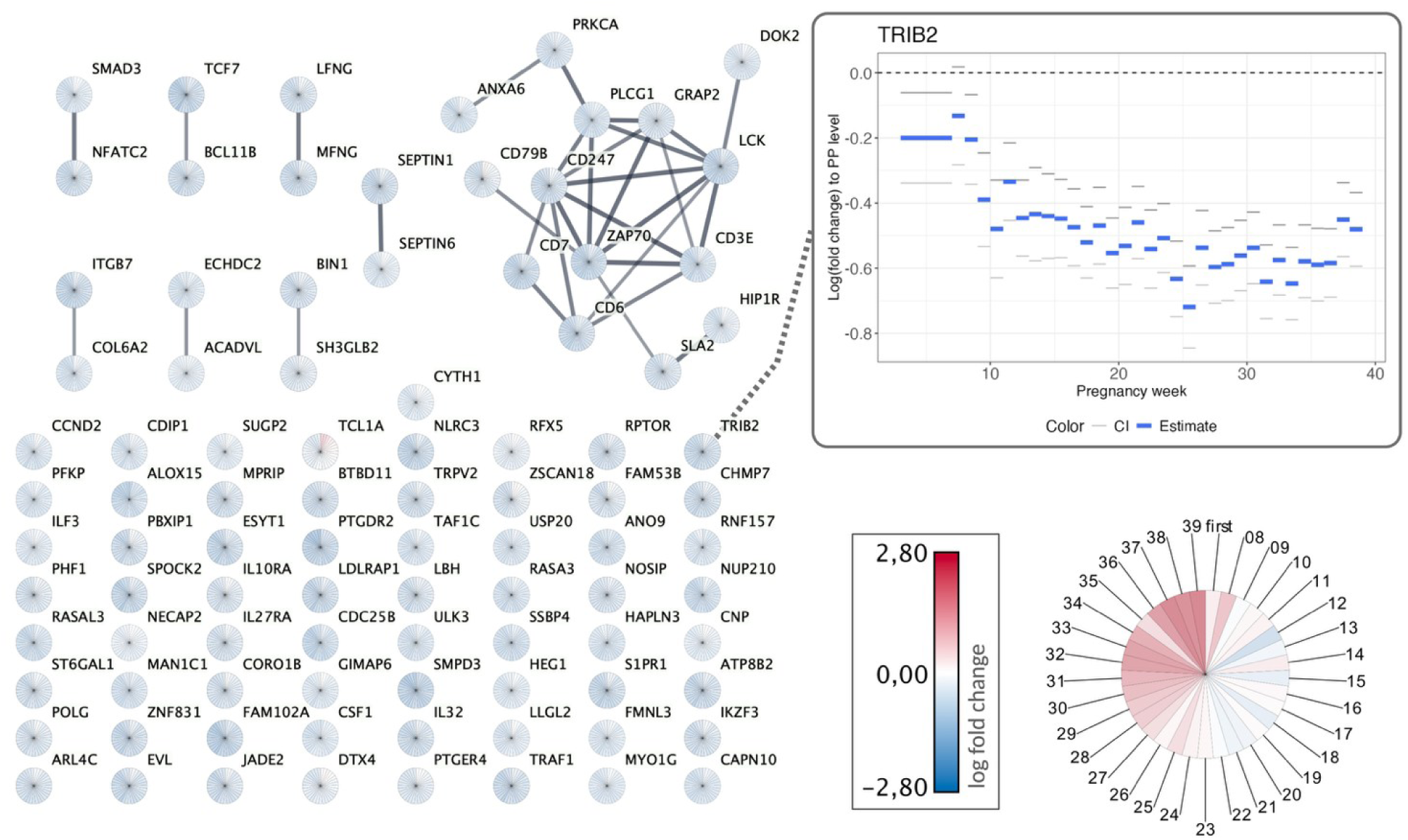
Functional associations among genes co-expressed in the brown module. Nodes represent individual genes, and edges reflect STRING database interactions with high confidence (score ≥ 0.7). Each node is visualized as a pie chart, where each segment corresponds to a pregnancy week from gestational weeks 4-7 (first), then to 8 and up to 39. The shading within each segment reflects the log fold change estimates in the corresponding week, mapped using a continuous blue to red color gradient for low to high expression. This radial representation highlights the temporal dynamics of gene expression across pregnancy at single-gene resolution. The highlighted node represents *TRIB2*, the module’s top gene in the discovery analysis. The plot shows the log fold change during pregnancy compared to the postpartum level with 95% confidence intervals.

### Modules with patterns of general, relatively stable higher expression levels

The turquoise and blue modules displayed expression levels that were higher than the postpartum reference levels and relatively stable across pregnancy (**Fig. 3**). Both modules were related to immune response.

The turquoise module (**Extended Data Fig. 6**) was enriched for terms associated with innate immune responses, including inflammation, neutrophil degranulation, cytokine signaling, and the Toll-like receptor (TLR) cascade (**Fig. 4**). Functional protein association analysis highlighted a network centered on *TLR1*, *TLR4*, *TLR8*, *TLR10* as well as with *S100A8*, *S100A9*, *S100A12*, and *LY96* (**Extended Data Fig. 6**).

The blue module (**Extended Data Fig. 7**) was enriched for terms related to immune response regulation, such as cytokine signaling pathways, and immune receptor activity, and was also associated with granules and vesicles (**Fig. 4**). Functional protein association analysis showed connections between *IL1B*, *IL1R1*, *IL1R2*, *IL1RA*, *CASP4*, *CASP5*, *TLR5*, *TLR6* and many more genes with roles in, e.g., interleukin-1 signaling and inflammasome activation (**Extended Data Fig. 7**).

### Changes in estimated cell-type proportions: pronounced increase in neutrophils with mid-pregnancy peak

To estimate the proportions of immune cell types during pregnancy, we performed deconvolution of gene-level expression data using CIBERSORTx with the LM22 signature matrix^17^. Seven cell types showed significant variation in proportions during pregnancy (**Supplementary Table 4**), including memory B cells, CD8 T cells, naïve and resting memory CD4 T cells, resting NK cells, monocytes, and neutrophils. Of these, neutrophils were the most prominent with increased proportions through most of pregnancy, particularly during mid- to late gestation (**Extended Data Fig. 8, Supplementary Fig. 3**), while memory B cells and CD8 T cells generally had the lowest levels. Memory B cells, monocytes, resting NK cells, and memory resting CD4 cells displayed relatively stable levels. Naïve CD4 T cells and CD8 T cells showed clear patterns of lower levels during pregnancy compared to the postpartum samples (**Extended Data Fig. 8, Supplementary Fig. 3**).

## Discussion

We present a high-resolution transcriptome-wide atlas of gene expression dynamics in maternal blood throughout healthy human pregnancy. Understanding the intricate biological processes of pregnancy requires longitudinal resolution beyond that afforded by conventional cross-sectional study designs. Our ultra-dense sampling – capturing around 25 time points per participant – offers an unprecedented view of the dynamic changes of the transcriptome across gestation. Using longitudinal modelling and a strict discovery/replication framework, we identified 720 genes with reproducible pregnancy-associated expression patterns. Co-expression network analysis organized these transcripts in nine modules with distinct temporal patterns – displaying gradual increases, gradual decreases, or general, relatively stable increased expression relative to the postpartum state. Functional enrichment and protein association analyses converged on numerous biological processes and pathways that included neutrophil-mediated immunity, activation of the innate immune response, antiviral responses, T cell differentiation and signaling, erythropoiesis and hemoglobin metabolism. Cell-type deconvolution analysis indicated changes in the proportions of seven cell types during pregnancy compared to the postpartum sample levels, including increased neutrophil levels and reduced levels of naïve CD4 and CD8 T cells.

Our findings reflect the complexities of maternal immune adaptation to simultaneously tolerate the semi-allogeneic fetus, modulate the different immunological stages of pregnancy, and maintain effective defense against pathogens^18^. In line with accumulating evidence of the importance of neutrophils in all stages of pregnancy from implantation and placentation through tissue remodeling to birth^19^, the black co-expression module – enriched for genes with roles in neutrophil degranulation and inflammatory signaling – showed progressively higher gene expression until mid-to-late pregnancy followed by a slight decline ahead of parturition. In contrast, the pink module displayed an initial decrease in expression in the first trimester followed by a sustained increase until delivery and was enriched for interferon-stimulated genes involved in the type I interferon antiviral response. This pattern aligns with evidence that type I interferon responses are essential for antiviral defense during pregnancy but that activation too early in gestation is associated with poor pregnancy outcomes, including poor placentation, intrauterine growth restriction, and fetal loss^20–22^. Furthermore, studies have shown associations between increased levels of Th1-derived cytokines, including interferon-γ, and miscarriage, small-for-gestational-age fetuses^23^, and reproductive failure^24^.

The role of T cells in pregnancy is often framed as a finely tuned shift toward Th2 and regulatory T cell (Treg) dominance that promotes maternal–fetal tolerance, accompanied by attenuation of Th1 and Th17 responses to prevent immunological rejection of the fetus^24–27^. Although this paradigm is most clearly observed at the maternal–fetal interface, maternal systemic shifts of T cell subsets have also been observed. For example, Th17 cell frequencies are reduced in healthy women in the third trimester of pregnancy compared with non-pregnant women^28^, and excessive Th17 responses have been associated with adverse outcomes such as recurrent miscarriage^24,26^. Consistent with this literature, the brown module was enriched for pathways involved in Th1, Th2, and Th17 cell differentiation. The pronounced decreasing expression of genes in this module may reflect a systemic transcriptional signature of immune adaptation, consistent with a reduced pro-inflammatory T cell profile and the promotion of tolerance as pregnancy progresses.

The metabolic demands of pregnancy require increases in plasma volume and red cell mass needed to meet increased circulatory system demands and ensure adequate oxygen delivery to the fetus^29^. Consistent with these physiological changes, the green co-expression module was enriched for pathways involved in hemopoiesis and erythrocyte differentiation and included many functional associations between genes related to erythropoiesis and heme biosynthesis.

The co-expression modules identified through WGCNA are inherently data-driven and unbiased, as they are constructed solely based on patterns of gene expression without relying on prior biological assumptions. In contrast, the functional enrichment analyses used to annotate these modules depend on existing literature and curated databases, which are often influenced by the extent to which specific biological areas have been studied^30,31^. The enrichment of immunological processes in multiple modules could thus reflect both the underlying biology and the comparatively high research attention given to immune pathways, including in the context of pregnancy^19,23,32^. Importantly, modules lacking strong enrichment for known biological functions should not be considered uninformative or unimportant; rather, they may capture transcriptional dynamics linked to biological processes that remain poorly characterized and therefore represent candidates for future investigation. We note, for example, that in all modules except the black module, the transcript with the lowest discovery *P* value was not connected to other proteins in the functional association analyses based on current literature, and several modules had only sparse overall connectivity.

Some of the observed changes in gene expression measured in whole blood could reflect changes in circulating cell-type composition during pregnancy rather than transcriptional regulation within cells. To address this possibility, we performed cell-type deconvolution analyses, which indicated significant differences in seven cell types, including increases in neutrophils and decreases in naïve CD4 and CD8 T cells. However, the estimated changes in cell type composition were less dynamic across pregnancy than the observed transcriptomic alterations, arguing that changes in cell type proportions alone are unlikely to account for the observed expression patterns. Importantly, the results should be interpreted with caution, as the LM22 signature matrix used for deconvolution was not derived from pregnant women, lacks temporal resolution, and only includes a subset of potentially relevant cell types. Future studies incorporating, e.g., pregnancy-specific reference profiles and/or single-cell RNA sequencing across gestation are warranted to illuminate the contribution of changes in cell-type proportions to global expression profiles during pregnancy.

Our study had limitations, including the lack of pre-pregnancy samples and limited coverage of the earliest stages of pregnancy. As a result, we may have missed some relevant transcriptional signals, e.g., associated with conception and implantation, during which critical immunological and endocrine processes occur^33^. Similarly, denser sampling closer to parturition would be required to fully capture the rapid transcriptional changes leading up to labor and delivery. Furthermore, as our study was conducted in a high-income setting, caution is warranted when generalizing findings to populations with different socioeconomic, demographic, environmental, or healthcare compositions. Despite the relatively limited number of participants, the dense longitudinal sampling scheme enabled robust modeling of gene expression dynamics over time. This design offers substantially finer temporal granularity compared to previous transcriptomic studies in pregnancy, which were constrained by cross-sectional designs or sparser sampling^6,7^. Notably, our use of a strict discovery/replication framework prioritized robustness of the findings, likely yielding a conservative number of transcripts with pregnancy-associated changes in expression. It is therefore plausible that many additional genes exhibit pregnancy-related expression changes, and we anticipate that pregnancy-associated patterns of transcription for many more of the >4,000 significant genes in the discovery analyses will be validated in future studies. To support further exploration, we provide results for all analyzed transcripts in an interactive interface that allows researchers to explore temporal patterns beyond the 720 robustly replicating genes reported here, and alongside it a possibility of further STRING network exploration. These open interfaces will support integrative analyses across additional cohorts and modalities.

In conclusion, our results provide unique insights into the finely tuned and tightly regulated transcriptional dynamics that support the progression of a healthy human pregnancy. We also establish a high-temporal-resolution reference atlas of maternal blood transcriptional dynamics across pregnancy to support future studies investigating disruptions in the molecular processes underlying pregnancy complications.

## Methods

### Study participants

The study was based on the Biological Signals in Pregnancy cohort established in Copenhagen, Denmark, between 2014 and 2019. To participate in the parent cohort, a woman had to be in the first trimester of pregnancy, free of chronic disease, and not using medication of any kind. For this study, a sub-cohort of 31 healthy pregnant women was randomly divided into a discovery cohort (n = 21) and a replication cohort (n = 10). The study was approved by the Scientific Ethics Committee for the Capital City Region of Denmark (H-3-2014-004) and Statens Serum Institut’s Department of Data Protection and Information Security, and reported to the Danish Data Protection Agency. All participants provided written informed consent.

### Sampling, RNA library preparation and RNA sequencing

Non-fasting blood samples were collected from all participants at weekly intervals from enrollment to delivery and once thereafter. Samples were collected in PaxGene Blood RNA tubes (BD Biosciences) and stored at -80°C until extraction. The samples from the discovery and replication cohorts were handled in separate batches. RNA was purified from whole blood using PaxGene Blood RNA kits (BD Biosciences). RNA quantity was assessed by NanoDrop (Thermo Fisher) and a subset of samples were checked for RNA integrity via BioAnalyzer (Agilent). Library preparation was performed using TruSeq Stranded Total RNA Preparation kits with Ribo-Zero Gold (Illumina) and assessed for library quantity by Qubit fluorometer (Thermo Fisher) and quality via Bioanalyzer. Samples were pooled at the per patient level and sequenced at 2×100 cycles on an Illumina HiSeq 4000 platform.

### Mapping, transcript quantification, cleaning and filtering

All reads from each sample were mapped with STAR v2.5.3a to the human reference genome GRCh38 (primary assembly) with GENCODE v28 (primary assembly) as transcript model reference. Gene-level quantification was obtained during the mapping by STAR (--quantMode TranscriptomeSAM GeneCounts).

A proxy for the degree of ribosomal depletion in each sample was created from the counts of all transcripts denoted as rRNA in GENCODE version 28 divided by the total number of mapped reads. The transcripts annotated to be rRNA were then removed from the data in both batches. The discovery and replication batch were filtered separately to exclude genes with low levels of expression. In the discovery batch a set of 32,104 detected transcripts was defined by requiring more than 0.5 counts per million (CPM) at a minimum of 3 time points in at least 75% of the participants, i.e. 15 out of 21 women in the discovery set. Likewise, 24,967 transcripts were detected in the replication batch by requiring more than 0.5 CPM at ≥3 time points in at least 7 of the 10 women in the replication set.

### Definition of adjusted pregnancy weeks

We defined pregnancy week as the unit of time measurement. Each pregnancy week was adjusted by multiplying the gestational age in days at the time of sampling by the ratio of a standard pregnancy length (280 days) and the actual length of the given pregnancy. This was done to collect transcriptional signals potentially associated with parturition around week 40 for all participants regardless of actual gestational week at delivery. The adjusted pregnancy week was used as the pregnancy week for a given sample. In the case of a participant delivering by scheduled Caesarean delivery, no gestational week adjustments were made. Furthermore, to avoid problems associated with sparse data in some gestational weeks, we considered all samples taken before week 8 in a single group. Similarly, because there was only a single sample in adjusted pregnancy week 40, this sample was analyzed together with samples from the previous week.

### Identification of pregnancy-associated transcripts

The 32,104 detected transcripts were tested for pregnancy-associated expression patterns by comparing expression levels during pregnancy with those measured in the postpartum samples using a log-linear mixed effects model (R version 4.5.0, lme4 package version 3.1-3^34^) in the discovery dataset. We use random effects to account for within-individual effects and fixed effects to correct for degree of ribosomal depletion, sequencing depth, and Trimmed Mean of M-values (TMM) normalization factors computed with edgeR^35^ utilizing default settings. We evaluated the difference in the gene expression counts at each adjusted pregnancy week relative to counts in the postpartum sample (reference level).

We validated the transcripts in the smaller replication cohort and then in the combined discovery and replication data using the same modeling setup. We used Bonferroni correction to determine the *P* value threshold for taking transcripts forward to the replication step and further required that the combined *P* value (discovery and replication sets combined) was lower than the *P* values in both the discovery and replication analysis alone, thereby defining a set of robustly replicated transcripts.

### Enrichment analysis of pregnancy-associated transcripts

We performed overrepresentation analysis of the set of pregnancy-associated transcripts compared to the statistical background of the whole genome using the STRING database’s web interface^8^. Enrichment analysis was conducted using STRING’s built-in functional enrichment tool, with default parameters, to perform hypergeometric tests to assess overrepresentation of GO terms^10^, KEGG^36^ and Reactome^11^ pathways, and other annotated sets. False discovery rate (FDR) correction, using default settings implemented in STRING^37^, was applied to control for multiple testing. Enriched terms were reported based on statistical significance (adjusted *P* value < 0.05), and biological relevance was assessed in the context of pregnancy-related processes.

### Gene-correlation networks

We applied co-expression analysis on the gene counts of the samples included in the discovery data set. Raw count of reads overlapping the quantified genes were normalized with variance stabilization transformation (VST) utilizing the DESeq package^38^. A further step of normalization was applied by regressing out the effect obtained from correction for ribosomal depletion, sequencing depth, and TMM normalization (see **Supplementary Fig. 4** for an illustration of the effects of the normalization steps). The residuals of the linear models were used as input for WGCNA^13^ to cluster the robustly replicated transcripts into co-expression modules (see details of the analysis in **Supplementary Fig 5**).

### STRING functional protein association network analysis

To explore potential associations among genes robustly associated with pregnancy and grouped into co-expression modules by WGCNA, we generated networks using functional protein association data from the STRING database^8^ (v12) and visualized them in Cytoscape^39^ (v3.10.2) through the RCy3 R package interface^40^. For each module, a list of gene identifiers was submitted to the Cytoscape stringapp^41^ (v2.2.0) via the string protein query command, using a minimum interaction confidence score of 0.7 (high confidence interactions), species restricted to *Homo sapiens* (NCBI Taxon ID: 9606), and excluding viral proteins.

The log-linear mixed model effect size (estimate) for a transcript of interest was mapped to the networks using the Omics Visualizer Cytoscape app^42^. Effect sizes were visualized with a blue-white-red gradient (range: −2.8 to 2.8). The layout of each STRING network was optimized using the Yfiles organic layout algorithm^43^ provided in Cytoscape.

### Cell-type deconvolution analysis

Gene counts for all genes were CPM-normalized with edgeR^35^. We deconvolved the gene level data with the docker version of CIBERSORTx^44^ with their accompanying LM22 signature matrix, where we applied B-mode (bulk mode) batch correction to remove technical differences between our dataset and the LM22 reference data profiled with different platforms (RNAseq and microarray, respectively). The deconvolution was performed in the default mode, thus obtaining relative cell-type proportions. The resulting proportions of 22 cell types were included in a linear mixed effects model with gestational week accounting for the effects of individual variation in cell type proportions using the proportions from the postpartum samples as the reference.

## Data availability

We implemented a Shiny app for users to be able to interactively explore the dynamic change in gene expression over pregnancy for all analyzed transcripts. It is available at https://github.com/ssi-dk/pregnancy-transcriptome.

We have also made interactive protein-protein networks available for the full set of 720 pregnancy-associated transcripts and for the nine WGCNA modules at the online STRING database (links in **Extended Data Table 1**). Individual-level RNA sequencing data from study participants are protected under the Danish Data Protection Act and the General Data Protection Regulation (GDPR), and therefore cannot be made publicly available.

## Code availability

Analysis code is available from https://github.com/ssi-dk/pregnancy-transcriptome.

## Acknowledgements

We are immensely grateful to the dedicated women who participated in the study, generously donating a blood sample every week during their pregnancy. We would also like to thank everyone involved in data collection and biological material handling.

B.F. received support from Independent Research Fund Denmark (0134-00244B). L.S. received support from a Carlsberg Foundation postdoctoral fellowship (CF15-0899). Partial funding was also obtained from the Oak Foundation. H.A.B was supported by the Novo Nordisk Foundation (NNF19OC0054286).

## Author Contributions

B.F., F.R.D.H., B.D.P., L.S., H.A.B., M.S., and M.M. conceptualized and designed the study with input from the remaining authors. M.-L.H.R., N.M.S., and M.M. organized and contributed to the collection of pregnancy samples. B.D.P. designed laboratory protocols, processed and analyzed the samples, with additional contributions and analytical input from L.L., C. J., F. V., E. W., Q. L., and H. C. F.R.D.H., L.S., K.N., J.P.S.F. and B.F. analyzed the data. H.A.B., M.S., and M.M. jointly supervised the study. B.F. and F.R.D.H. wrote the first manuscript draft, and B.D.P., L.S., K.N., L.L., J.P.S.F., M.-L.H.R., N.M.S., C. J., F. V., E. W., Q. L., H. C., F.G., H.A.B., M.S., and M.M. contributed with interpretation of results and reviewing and editing the manuscript.

## Competing interests

L.L. is a co-founder of NiMo Therapeutics. J.F. is an employee of Novo Nordisk. M.P.S. is a cofounder and scientific advisor of Crosshair Therapeutics, Exposomics, Filtricine, Fodsel, Iollo, InVu Health, January AI, Marble Therapeutics, Mirvie, Next Thought AI, Orange Street Ventures, Personalis, Protos Biologics, Qbio, RTHM, and SensOmics. He is also a scientific advisor of Abbratech, Applied Cognition, Enovone, Jupiter Therapeutics, M3 Helium, Mitrix, Neuvivo, Onza, Sigil Biosciences, TranscribeGlass, WndrHLTH, and Yuvan Research. M.P.S. is a co-founder of NiMo Therapeutics. He is an investor and scientific advisor of R42 and Swaza and also an investor in Repair Biotechnologies. M.M. is a cofounder and scientific advisor of Mirvie.

## Extended Data

**Extended Data Fig. 1:**
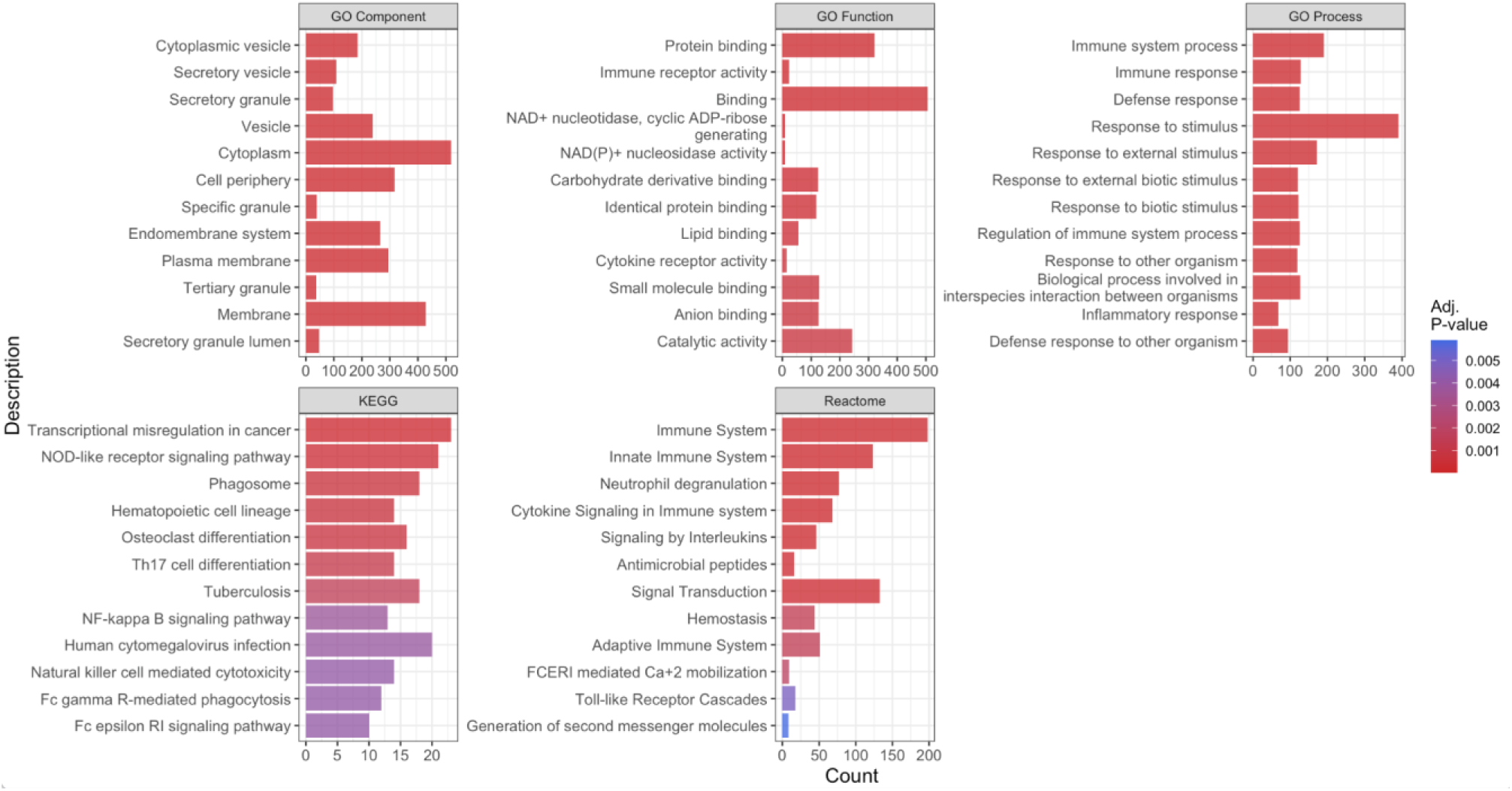
Top 12 enriched terms for the set of transcripts associated with pregnancy. Bar plots display the most significantly enriched Gene Ontology Biological Processes, Molecular Functions, Cellular Components, KEGG, and Reactome pathways. Enrichment was assessed using the STRING database, with count representing the number of genes enriched for the given terms, and color the Benjamini-Hochberg adjusted *P* values for each term.

**Extended Data Fig. 2:**
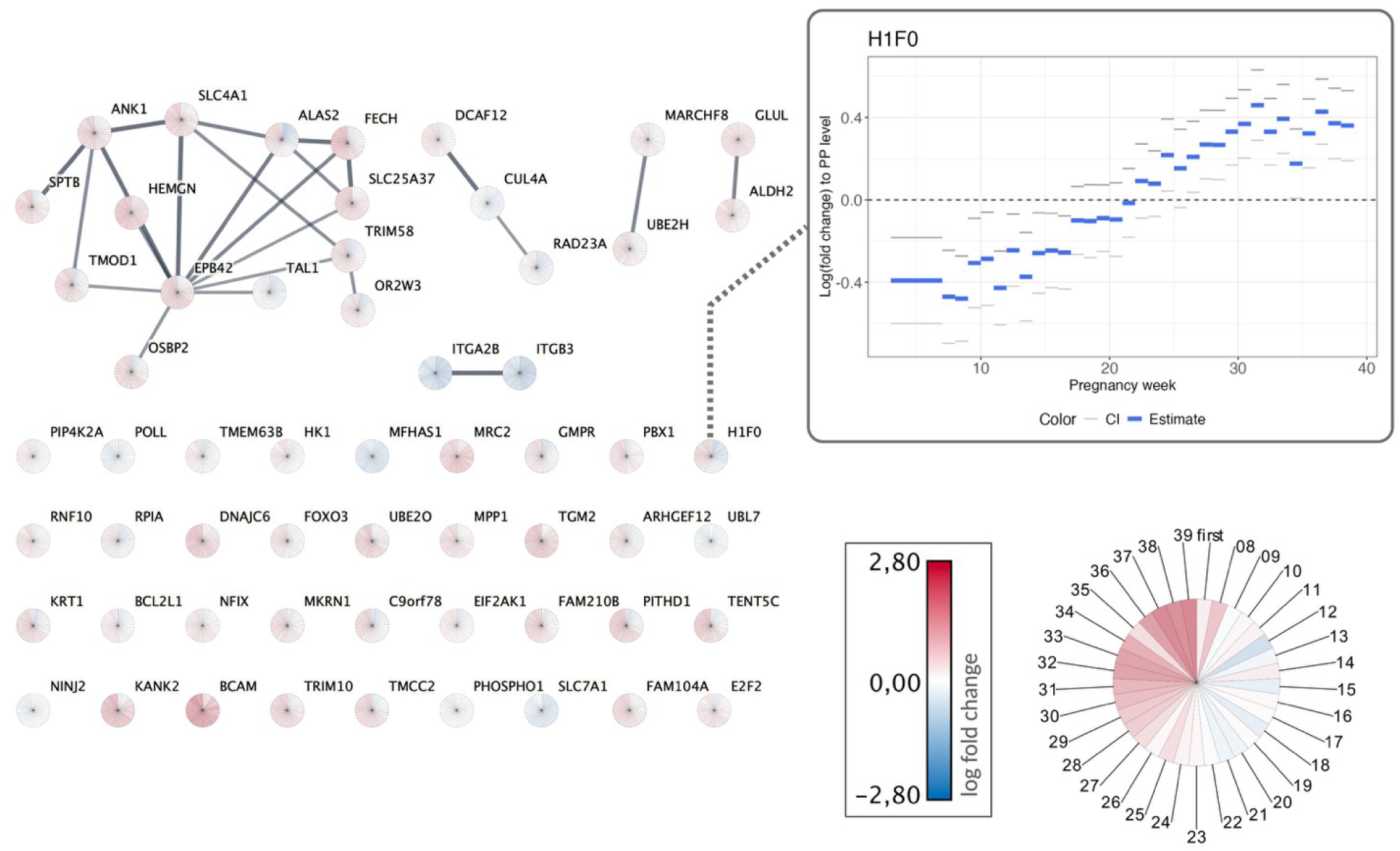
Functional associations among genes co-expressed in the green module. Nodes represent individual genes, and edges reflect STRING database interactions with high confidence (score ≥ 0.7). Each node is visualized as a pie chart, where each segment corresponds to a pregnancy week from gestational weeks 4-7 (first), then to 8 and up to 39. The shading within each segment reflects gene expression estimates in the corresponding week, mapped using a continuous blue to red color gradient for low to high expression. The highlighted node represents *H1F0*, the module’s top gene in the discovery analysis. The plot shows the log fold change during pregnancy compared to the postpartum level with 95% confidence intervals.

**Extended Data Fig. 3:**
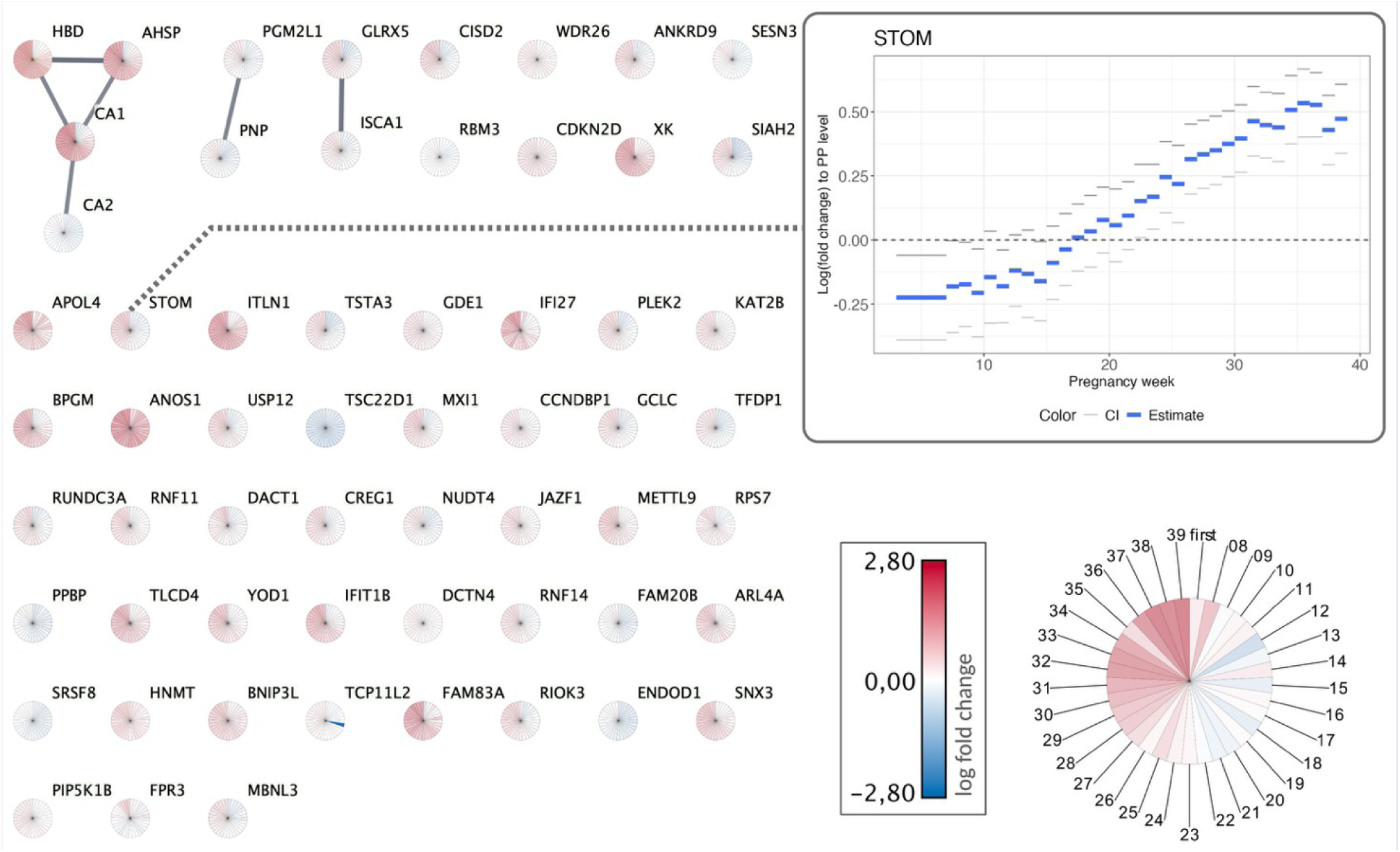
Functional associations among genes co-expressed in the yellow module. Nodes represent individual genes, and edges reflect STRING database interactions with high confidence (score ≥ 0.7). Each node is visualized as a pie chart, where each segment corresponds to a pregnancy week from gestational weeks 4-7 (first), then to 8 and up to 39. The shading within each segment reflects gene expression estimates in the corresponding week, mapped using a continuous blue to red color gradient for low to high expression. The highlighted node represents *STOM*, the module’s top gene in the discovery analysis. The plot shows the log fold change during pregnancy compared to the postpartum level with 95% confidence intervals.

**Extended Data Fig. 4:**
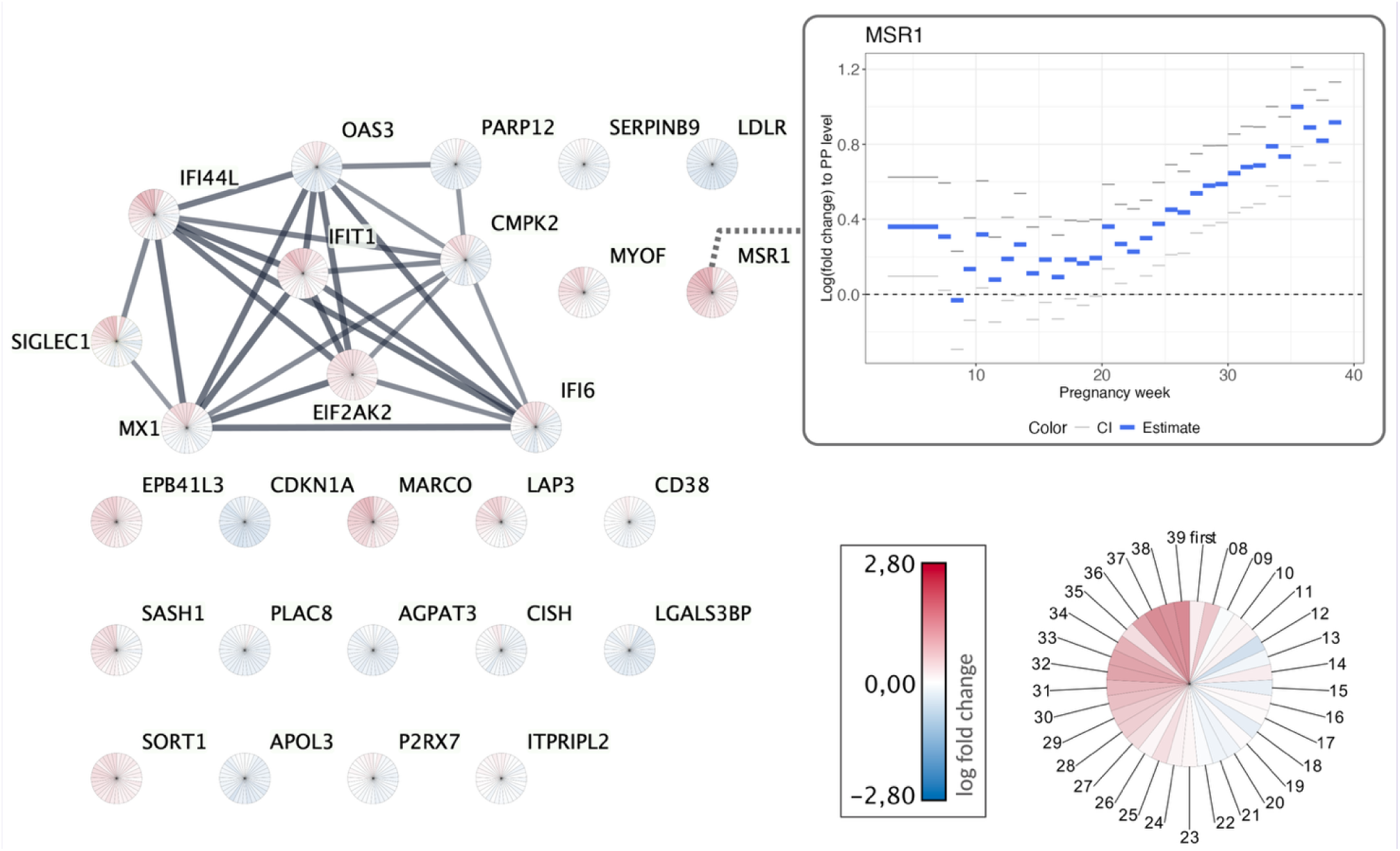
Functional associations among genes co-expressed in the pink module. Nodes represent individual genes, and edges reflect STRING database interactions with high confidence (score ≥ 0.7). Each node is visualized as a pie chart, where each segment corresponds to a pregnancy week from gestational week 4-7 (first), then to 8 and up to 39. The shading within each segment reflects gene expression estimates in the corresponding week, mapped using a continuous blue to red color gradient for low to high expression. The highlighted node represents *MSR1*, the module’s top gene in the discovery analysis. The plot shows the log fold change during pregnancy compared to the postpartum level with 95% confidence intervals.

**Extended Data Fig. 5:**
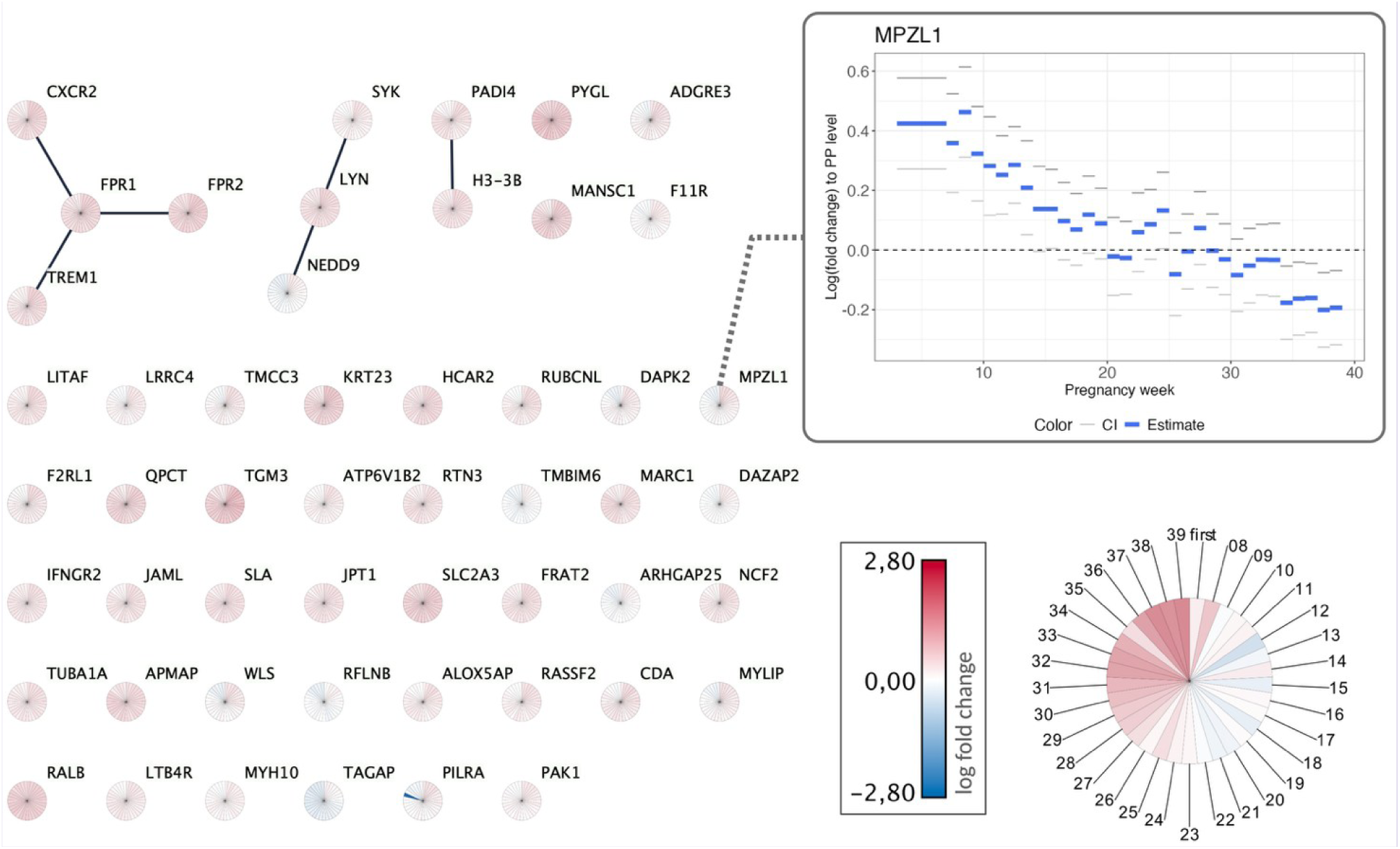
Functional associations among genes co-expressed in the red module. Nodes represent individual genes, and edges reflect STRING database interactions with high confidence (score ≥ 0.7). Each node is visualized as a pie chart, where each segment corresponds to a pregnancy week from gestational week 4-7 (first), then to 8 and up to 39. The shading within each segment reflects gene expression estimates in the corresponding week, mapped using a continuous blue to red color gradient for low to high expression. The highlighted node represents *MPZL1*, the module’s top gene in the discovery analysis. The plot shows the log fold change during pregnancy compared to the postpartum level with 95% confidence intervals.

**Extended Data Fig. 6:**
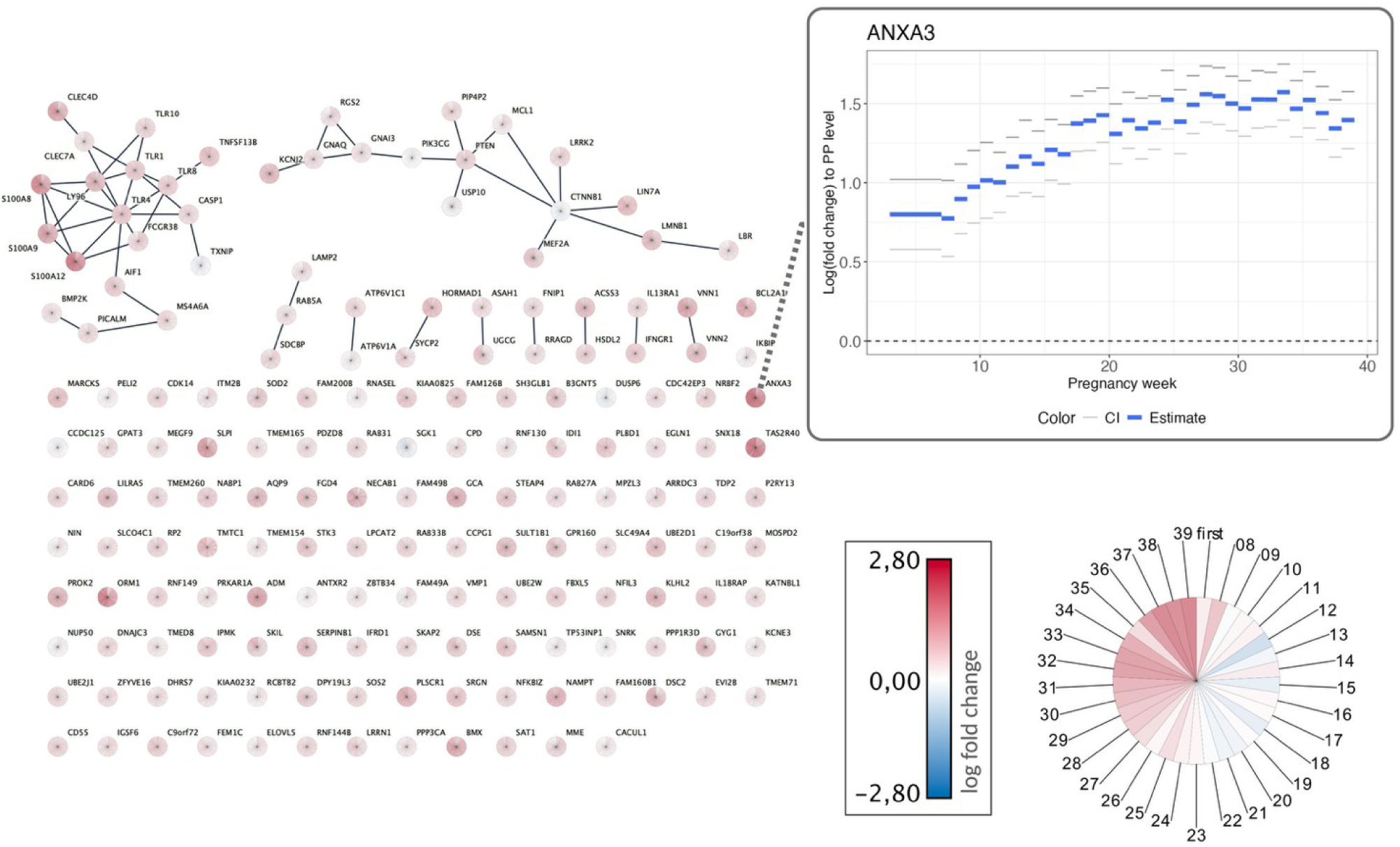
Functional associations among genes co-expressed in the turquoise module. Nodes represent individual genes, and edges reflect STRING database interactions with high confidence (score ≥ 0.7). Each node is visualized as a pie chart, where each segment corresponds to a pregnancy week from gestational week 4-7 (first), then to 8 and up to 39. The shading within each segment reflects gene expression estimates in the corresponding week, mapped using a continuous blue to red color gradient for low to high expression. The highlighted node represents *ANXA3*, the module’s top gene in the discovery analysis. The plot shows the log fold change during pregnancy compared to the postpartum level with 95% confidence intervals.

**Extended Data Fig. 7:**
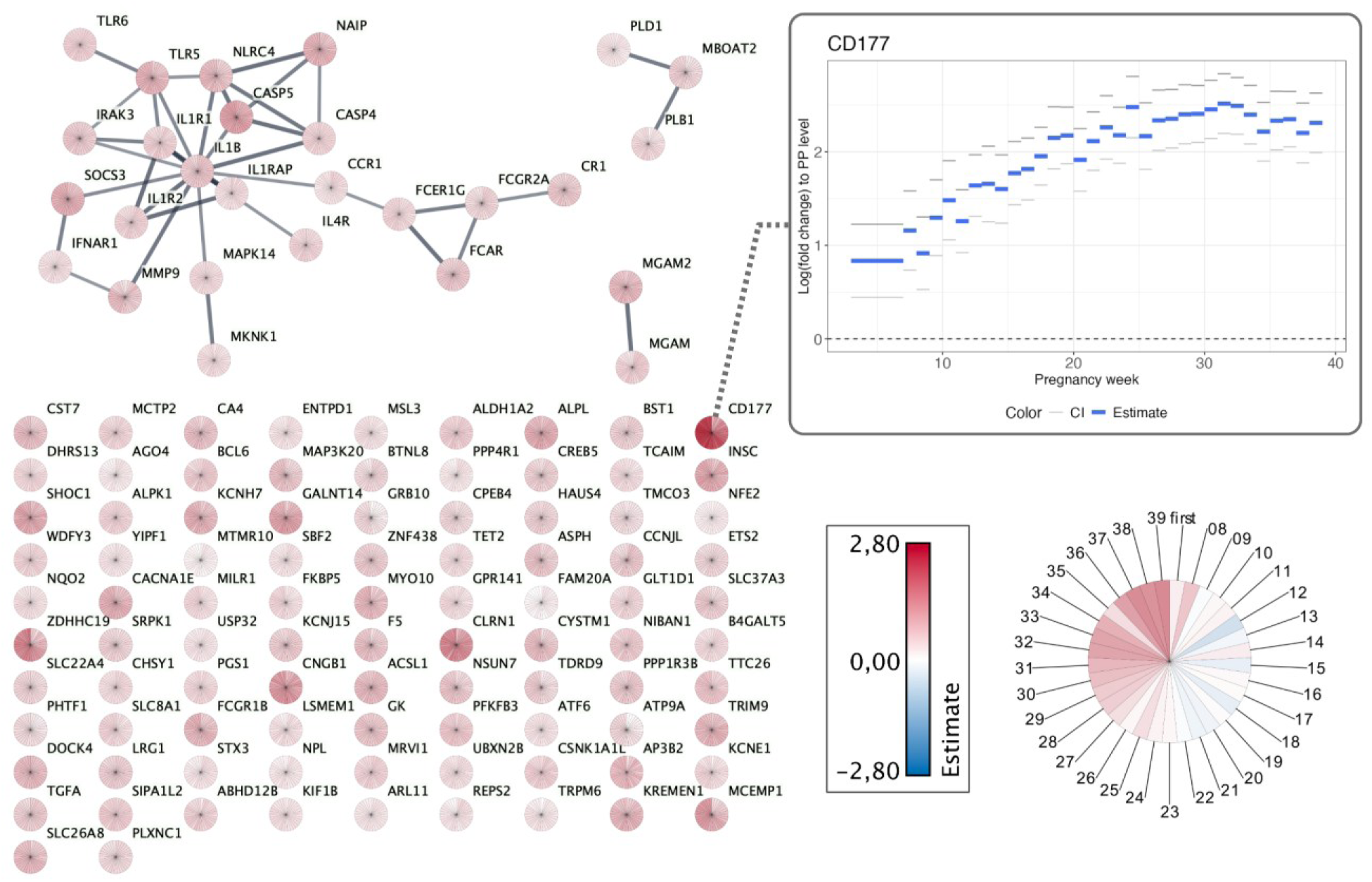
Functional associations among genes co-expressed in the blue module. Nodes represent individual genes, and edges reflect STRING database interactions with high confidence (score ≥ 0.7). Each node is visualized as a pie chart, where each segment corresponds to a pregnancy week from gestational week 4-7 (first), then to 8 and up to 39. The shading within each segment reflects the log fold change estimates in the corresponding week, mapped using a continuous blue to red color gradient for low to high expression. The highlighted node represents *CD177*, the module’s top gene in the discovery analysis. The plot shows the log fold change during pregnancy compared to the postpartum level with 95% confidence intervals.

**Extended Data Fig. 8:**
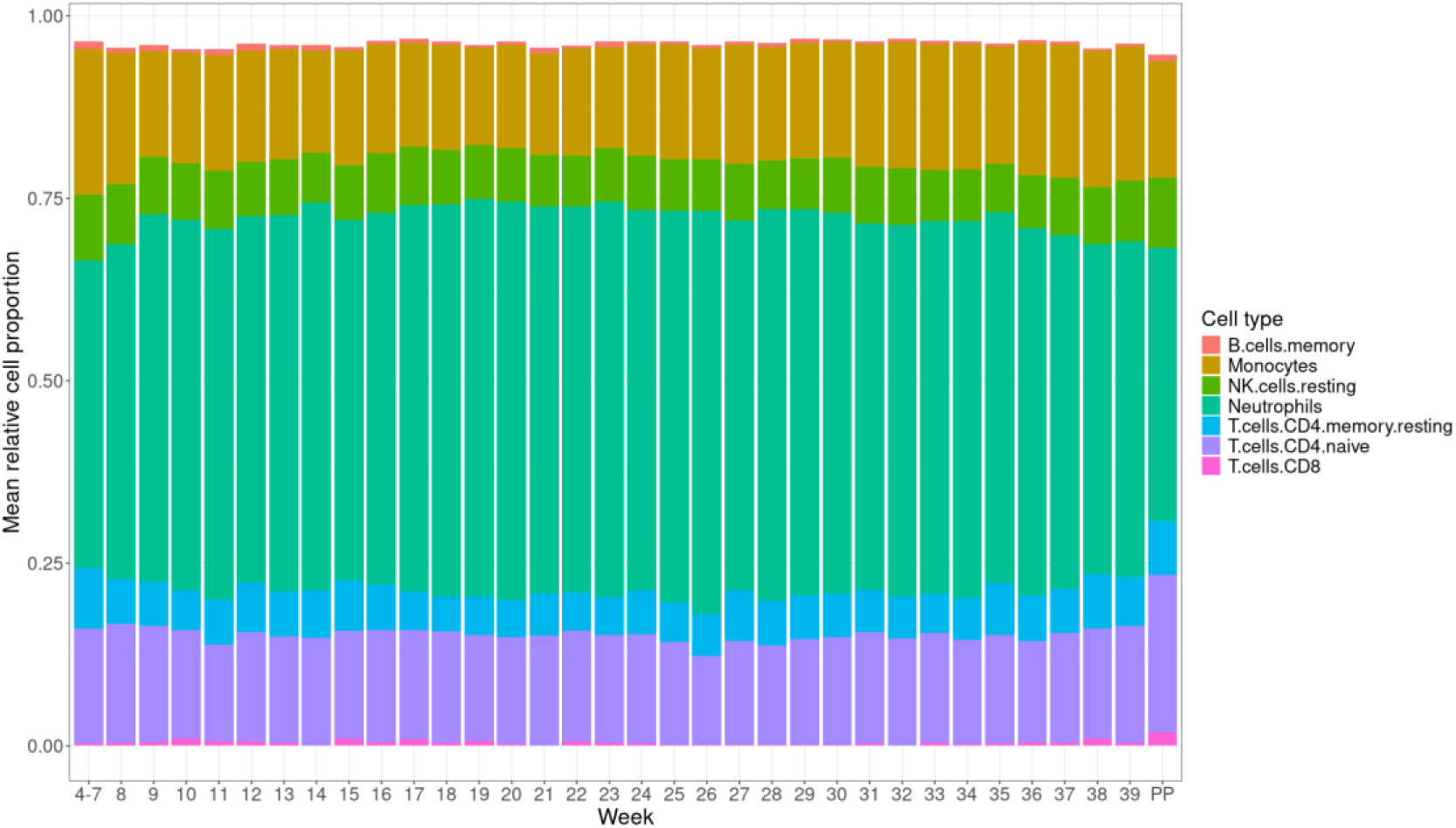
Stacked bar plot of estimated cell-type proportions. Results for the seven cell types that showed significantly different proportions during pregnancy compared to the postpartum samples. Shown are all sample cell proportions grouped for all measured pregnancy weeks.

**Extended Data Table 1:** Permanent links to STRING enrichment analyses. Results are given for the full set of pregnancy-associated transcripts and for each of the nine WGCNA derived co-expression modules. For each protein set, we report the protein-protein interaction (PPI) enrichment *P* value, which tests whether the proteins show more interaction among themselves than expected for a random set of proteins with the same size network, indicating non-random biological connectivity. The networks can be accessed through the links and explored interactively, and the different tables can be downloaded.

| Set of genes | PPI Enrichment | STRING Permalinks |
| --- | --- | --- |
| Full set of pregnancy associated transcripts | $P < 1.0 \times 10^{-16}$ | https://version-12-0.string-db.org/cgi/network?networkId=bJijKJ92QKrG |
| Green module | $P = 3.51 \times 10^{-14}$ | https://version-12-0.string-db.org/cgi/network?networkId=bdZxFhctUVUh |
| Yellow module | $P = 0.0142$ | https://version-12-0.string-db.org/cgi/network?networkId=bd26XylFkaIE |
| Black module | $P < 1.0 \times 10^{-16}$ | https://version-12-0.string-db.org/cgi/network?networkId=bK9hakaRQyX2 |
| Pink module | $P < 1.0 \times 10^{-16}$ | https://version-12-0.string-db.org/cgi/network?networkId=bGasjihl31aH |
| Red module | 0.0247 | https://version-12-0.string-db.org/cgi/network?networkId=byv2VvUpbr7i |
| Brown module | $P = 4.14 \times 10^{-12}$ | https://version-12-0.string-db.org/cgi/network?networkId=bjQnlx0Y3ItK |
| Turquoise module | $P = 2.74 \times 10^{-8}$ | https://version-12-0.string-db.org/cgi/network?networkId=bGGmXuwfrYd2 |
| Blue module | $P = 3.18 \times 10^{-12}$ | https://version-12-0.string-db.org/cgi/network?networkId=bWWbpid2tM3I |
| Grey module | $P = 3.71 \times 10^{-13}$ | https://version-12-0.string-db.org/cgi/network?networkId=bf1ciayjA24c |

## References

1. Hod, T., Cerdeira, A. S. & Karumanchi, S. A. Molecular mechanisms of preeclampsia. Cold Spring Harb. Perspect. Med. 5, a023473 (2015).

2. Cuffe, J. S. M., Holland, O., Salomon, C., Rice, G. E. & Perkins, A. V. Placental derived biomarkers of pregnancy disorders. Placenta 54, 104–110 (2017).

3. Fejzo, M. S. et al. Placenta and appetite genes GDF15 and IGFBP7 are associated with hyperemesis gravidarum. Nat. Commun. 9, 1178 (2018).

4. Jain, V. G., Monangi, N., Zhang, G. & Muglia, L. J. Genetics, epigenetics, and transcriptomics of preterm birth. Am. J. Reprod. Immunol. 88, e13600 (2022).

5. Elovitz, M. A. et al. Molecular subtyping of hypertensive disorders of pregnancy. Nat. Commun. 16, 1–14 (2025).

6. Gomez-Lopez, N. et al. The cellular transcriptome in the maternal circulation during normal pregnancy: a longitudinal study. Front. Immunol. 10, 2863 (2019).

7. Wright, M. L. et al. Pregnancy-associated systemic gene expression compared to a pre-pregnancy baseline, among healthy women with term pregnancies. Front. Immunol. 14, 1161084 (2023).

8. Szklarczyk, D. et al. The STRING database in 2023: protein--protein association networks and functional enrichment analyses for any sequenced genome of interest. Nucleic Acids Res. 51, D638--D646 (2023).

9. Dyer, S. C. et al. Ensembl 2025. Nucleic Acids Res. 53, D948--D957 (2025).

10. Aleksander, S. A. et al. The gene ontology knowledgebase in 2023. Genetics 224, iyad031 (2023).

11. Milacic, M. et al. The reactome pathway knowledgebase 2024. Nucleic Acids Res. 52, D672--D678 (2024).

12. Kanehisa, M. & Goto, S. KEGG: kyoto encyclopedia of genes and genomes. Nucleic Acids Res. 28, 27–30 (2000).

13. Langfelder, P. & Horvath, S. WGCNA: an R package for weighted correlation network analysis. BMC Bioinformatics 9, 559 (2008).

14. Rørvig, S., Østergaard, O., Heegaard, N. H. H. & Borregaard, N. Proteome profiling of human neutrophil granule subsets, secretory vesicles, and cell membrane: correlation with transcriptome profiling of neutrophil precursors. J. Leukoc. Biol. 94, 711–721 (2013).

15. Rahkonen, L. et al. Factors affecting matrix metalloproteinase-8 levels in the vaginal and cervical fluids in the first and second trimester of pregnancy. Hum. Reprod. 24, 2693–2702 (2009).

16. Rørvig, S. et al. Ficolin-1 is present in a highly mobilizable subset of human neutrophil granules and associates with the cell surface after stimulation with fMLP. J. Leukoc. Biol. 86, 1439–1449 (2009).

17. Newman, A. M. et al. Determining cell type abundance and expression from bulk tissues with digital cytometry. Nat. Biotechnol. 37, 773–782 (2019).

18. Mor, G., Aldo, P. & Alvero, A. B. The unique immunological and microbial aspects of pregnancy. Nat. Rev. Immunol. 17, 469–482 (2017).

19. Bert, S., Ward, E. J. & Nadkarni, S. Neutrophils in pregnancy: New insights into innate and adaptive immune regulation. Immunology 164, 665–676 (2021).

20. Casazza, R. L., Lazear, H. M. & Miner, J. J. Protective and pathogenic effects of interferon signaling during pregnancy. Viral Immunol. 33, 3–11 (2020).

21. Buchrieser, J. et al. IFITM proteins inhibit placental syncytiotrophoblast formation and promote fetal demise. Science (80-). 365, 176–180 (2019).

22. Simoni, M. K. et al. Type I interferon exposure of an implantation-on-a-chip device alters invasive extravillous trophoblast function. Cell Reports Med. 6, (2025).

23. Weetman, A. P. The immunology of pregnancy. Thyroid 9, 643–646 (1999).

24. La Rocca, C., Carbone, F., Longobardi, S. & Matarese, G. The immunology of pregnancy: regulatory T cells control maternal immune tolerance toward the fetus. Immunol. Lett. 162, 41–48 (2014).

25. Saito, S., Nakashima, A., Shima, T. & Ito, M. Th1/Th2/Th17 and regulatory T-cell paradigm in pregnancy. Am. J. Reprod. Immunol. 63, 601–610 (2010).

26. Wang, W., Sung, N., Gilman-Sachs, A. & Kwak-Kim, J. T helper (Th) cell profiles in pregnancy and recurrent pregnancy losses: Th1/Th2/Th9/Th17/Th22/Tfh cells. Front. Immunol. 11, 2025 (2020).

27. Figueiredo, A. S. & Schumacher, A. The T helper type 17/regulatory T cell paradigm in pregnancy. Immunology 148, 13–21 (2016).

28. Santner-Nanan, B. et al. Systemic increase in the ratio between Foxp3+ and IL-17-producing CD4+ T cells in healthy pregnancy but not in preeclampsia. J. Immunol. 183, 7023–7030 (2009).

29. Hytten, F. Blood volume changes in normal pregnancy. Clin. Haematol. 14, 601–612 (1985).

30. Stoeger, T., Gerlach, M., Morimoto, R. I. & Nunes Amaral, L. A. Large-scale investigation of the reasons why potentially important genes are ignored. PLoS Biol. 16, e2006643 (2018).

31. Kustatscher, G. et al. An open invitation to the understudied proteins initiative. Nat. Biotechnol. 40, 815–817 (2022).

32. Aghaeepour, N., et al. An immune clock of human pregnancy. Sci. Immunol. 2, eaan2946 (2017).

33. Van Mourik, M. S. M., Macklon, N. S. & Heijnen, C. J. Embryonic implantation: cytokines, adhesion molecules, and immune cells in establishing an implantation environment. J. Leucoc. Biol. 85, 4–19 (2009).

34. Bates, D., Mächler, M., Bolker, B. & Walker, S. Fitting linear mixed-effects models using lme4. J. Stat. Softw. 67, 1–48 (2015).

35. Chen, Y., Chen, L., Lun, A. T. L., Baldoni, P. L. & Smyth, G. K. edgeR v4: powerful differential analysis of sequencing data with expanded functionality and improved support for small counts and larger datasets. Nucleic Acids Res. 53, gkaf018 (2025).

36. Kanehisa, M., Furumichi, M., Sato, Y., Kawashima, M. & Ishiguro-Watanabe, M. KEGG for taxonomy-based analysis of pathways and genomes. Nucleic Acids Res. 51, D587--D592 (2023).

37. Szklarczyk, D. et al. The STRING database in 2025: protein networks with directionality of regulation. Nucleic Acids Res. 53, D730--D737 (2025).

38. Love, M. I., Huber, W. & Anders, S. Moderated estimation of fold change and dispersion for RNA-seq data with DESeq2. Genome Biol. 15, 1–21 (2014).

39. Shannon, P. et al. Cytoscape: a software environment for integrated models of biomolecular interaction networks. Genome Res. 13, 2498–2504 (2003).

40. Gustavsen, J. A., Pai, S., Isserlin, R., Demchak, B. & Pico, A. R. RCy3: Network biology using Cytoscape from within R. F1000Research 8, 1774 (2019).

41. Doncheva, N. T. et al. Cytoscape stringApp 2.0: analysis and visualization of heterogeneous biological networks. J. Proteome Res. 22, 637–646 (2022).

42. Legeay, M., Doncheva, N. T., Morris, J. H. & Jensen, L. J. Visualize omics data on networks with Omics Visualizer, a Cytoscape App. F1000Research 9, 157 (2020).

43. Wiese, R., Eiglsperger, M. & Kaufmann, M. yfiles—visualization and automatic layout of graphs. in Graph Drawing Software 173–191 (Springer, 2004).

44. Wang, X., Park, J., Susztak, K., Zhang, N. R. & Li, M. Bulk tissue cell type deconvolution with multi-subject single-cell expression reference. Nat. Commun. 10, 380 (2019).

