## Supplementary Information for "An atlas of transcriptional dynamics in maternal blood over the course of healthy pregnancy"

### Supplementary Text

#### Sequencing depth and mapping

The discovery samples were sequenced at an average of 29.1 million reads per sample. Of these reads 71.3% were uniquely mapped to the human genome (GRCh38 + GENCODE v28 primary assembly), while 22.9% were multi-mapping reads and 5.8% remained unmapped.

The replication samples were sequenced at an average of 20.6 million reads per sample and of these reads 50.9% were uniquely mapped to the human genome (GRCh38 + GENCODE v28 primary assembly), while 45.2% were multi-mapping reads and 3.9% remained unmapped

In total the mapping yielded 13,953 million uniquely mapped reads.

#### Quantification of expression levels for known features

The set of 50K known transcripts used for annotation while mapping included 19K protein coding transcripts, 17K processed transcripts and 12K pseudogenes (Ensembl biotypes). Reads overlapping these known transcripts were quantified from the uniquely mapped reads.

In the discovery samples, 70.2% of uniquely mapped reads were uniquely assigned to a single transcript, while 24.4% overlapped multiple features and 5.4% were not assigned to any feature. In the replication samples, 52.7% of mapped reads were uniquely assigned to a single transcript, while 45.2% overlapped multiple features and 2.1% were not assigned to any feature.

From reads overlapping known features, 551 transcripts annotated to be ribosomal RNA from GENCODE v28 primary assembly were removed from both the discovery and replication batches. Furthermore, an identification of repeatedly detected transcripts (>0.5 CPM at > 2 timepoints in 75% of sample batch women) resulted in 32104 transcripts and 24967 transcripts in the discovery and replication batch, respectively.

### Supplementary Figures

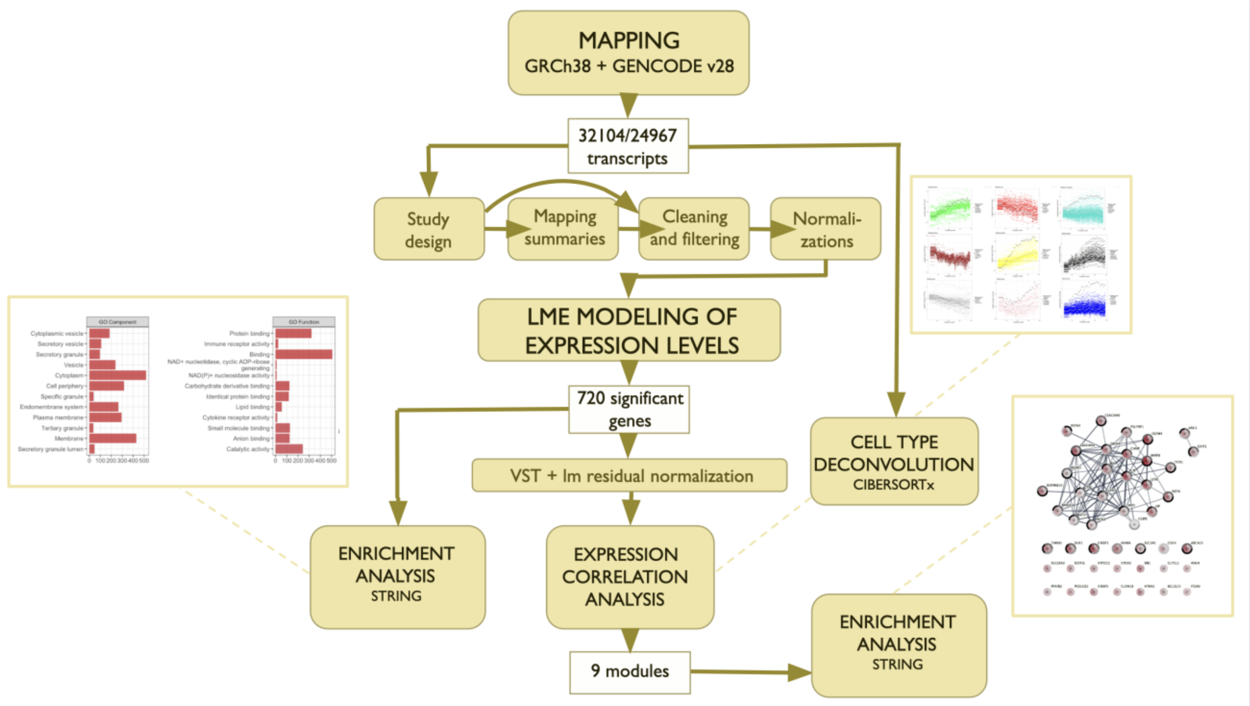

**Supplementary Fig. 1: Flow chart of the study design.**

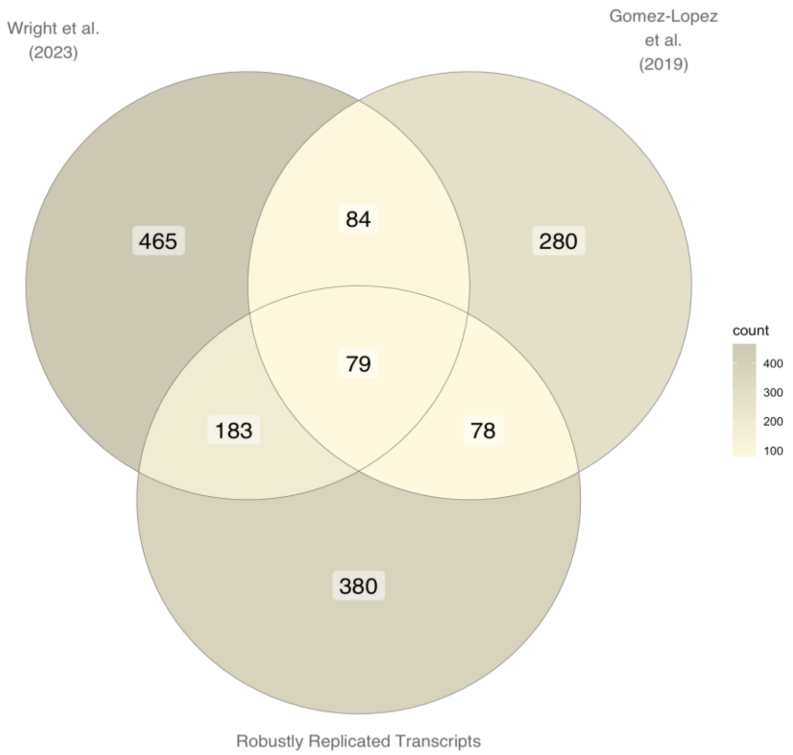

**Supplementary Fig. 2:** Venn diagram of overlap between identified pregnancy-related transcripts found in Wright et al. (2023) REF and REF Gomez-Lopes et al. (2019).

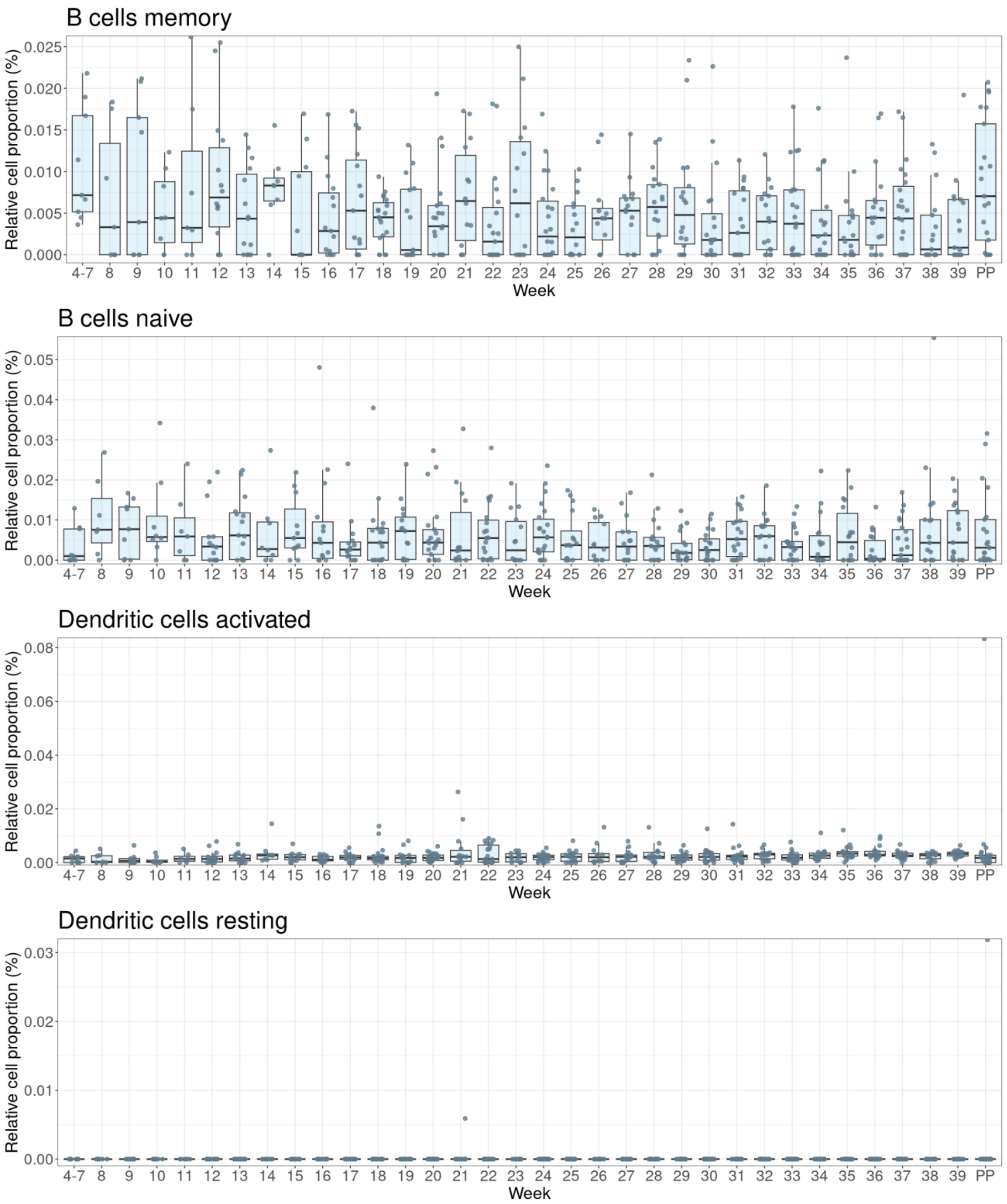

B cells memory
B cells naive
Dendritic cells activated
Dendritic cells resting
Relative cell proportion (%)
Week
4-7
PP

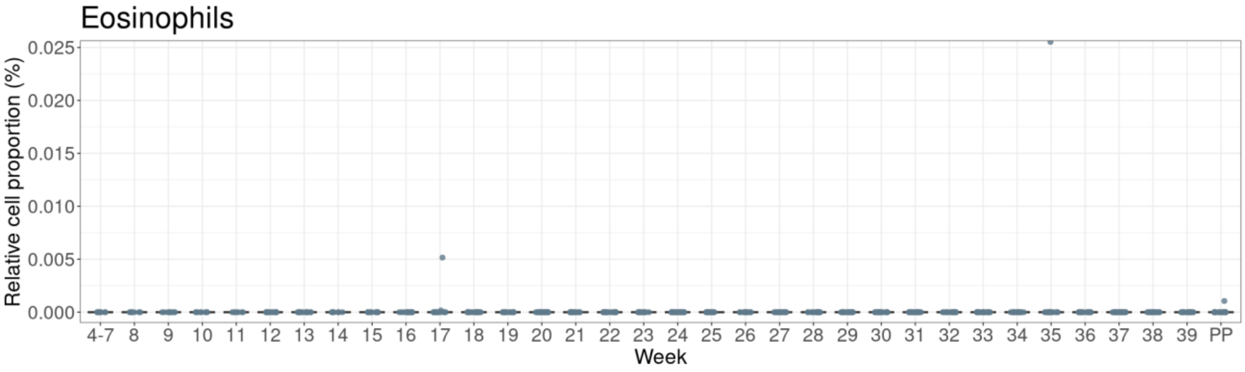
Eosinophils
Relative cell proportion (%)
0.025
0.020
0.015
0.010
0.005
0.000
4-7 8 9 10 11 12 13 14 15 16 17 18 19 20 21 22 23 24 25 26 27 28 29 30 31 32 33 34 35 36 37 38 39 PP
Week

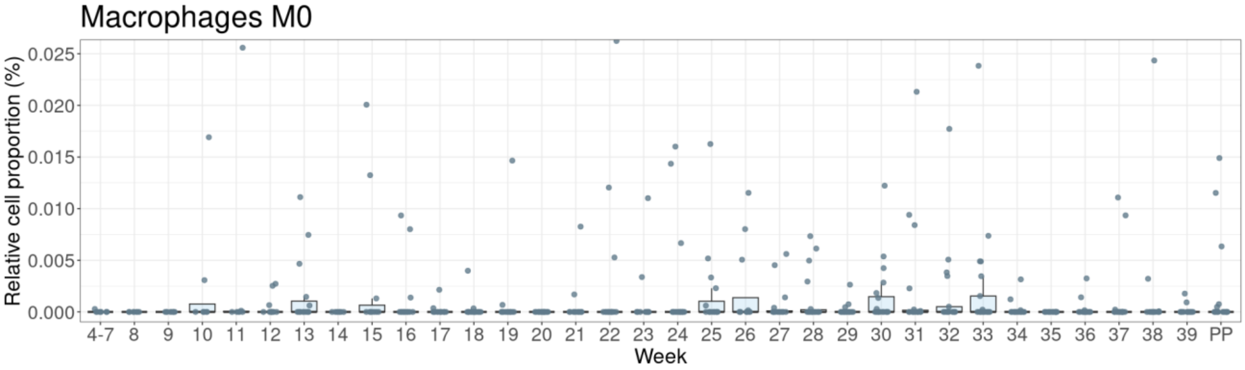
Macrophages M0
Relative cell proportion (%)
0.025
0.020
0.015
0.010
0.005
0.000
4-7 8 9 10 11 12 13 14 15 16 17 18 19 20 21 22 23 24 25 26 27 28 29 30 31 32 33 34 35 36 37 38 39 PP
Week

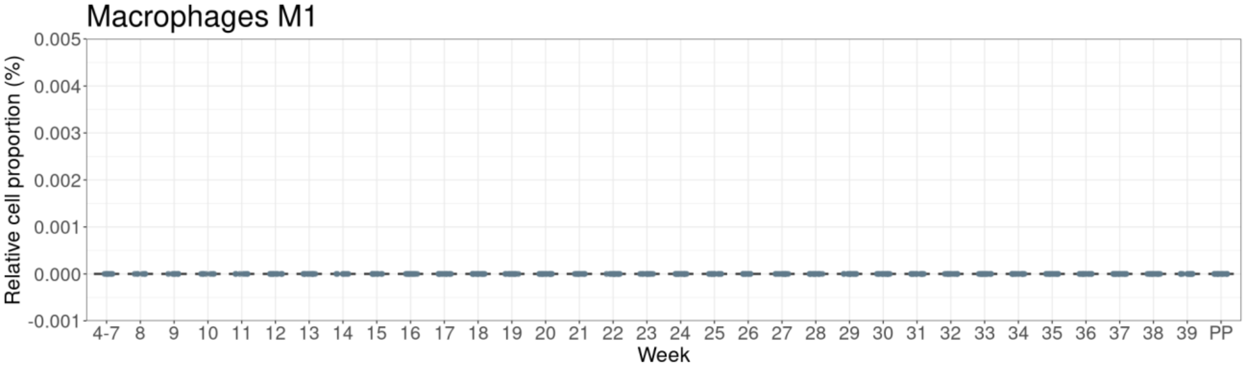
Macrophages M1
Relative cell proportion (%)
0.005
0.004
0.003
0.002
0.001
0.000
-0.001
4-7 8 9 10 11 12 13 14 15 16 17 18 19 20 21 22 23 24 25 26 27 28 29 30 31 32 33 34 35 36 37 38 39 PP
Week

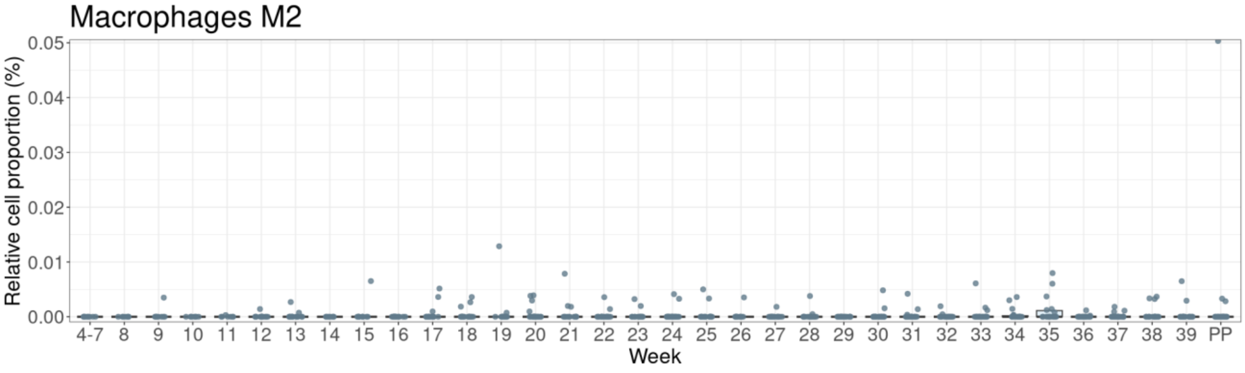
Macrophages M2
Relative cell proportion (%)
0.05
0.04
0.03
0.02
0.01
0.00
4-7 8 9 10 11 12 13 14 15 16 17 18 19 20 21 22 23 24 25 26 27 28 29 30 31 32 33 34 35 36 37 38 39 PP
Week

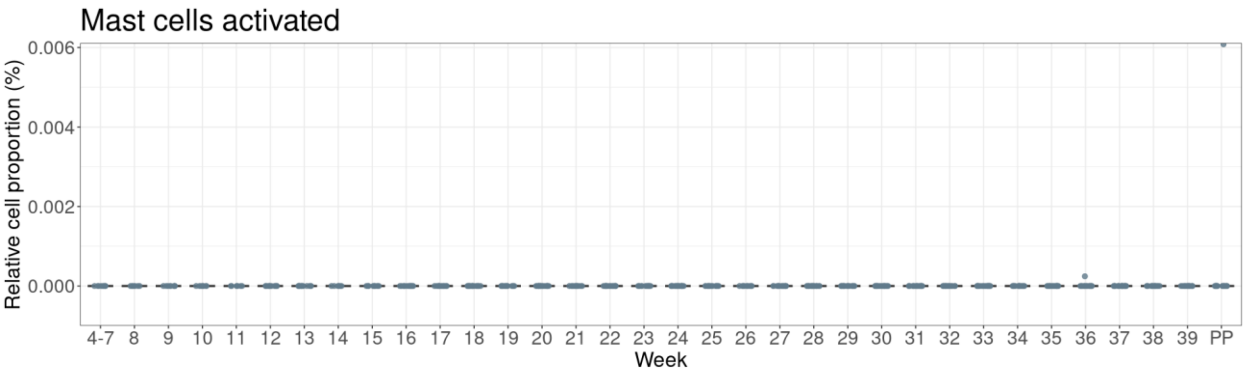
Mast cells activated
Relative cell proportion (%)
Week

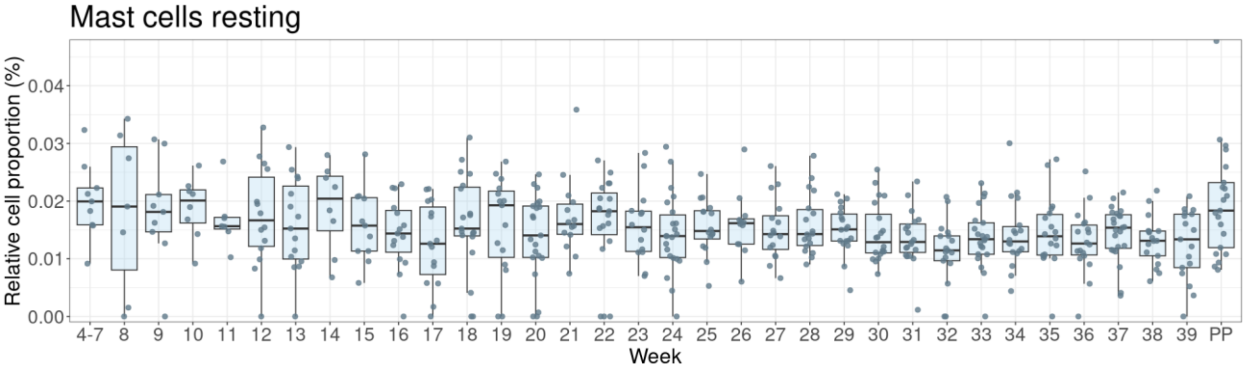
Mast cells resting
Relative cell proportion (%)
Week

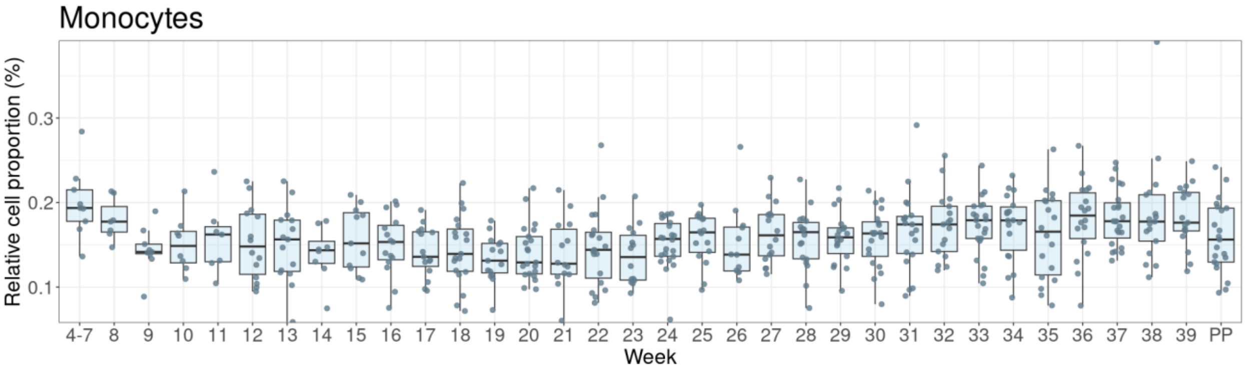
Monocytes
Relative cell proportion (%)
Week

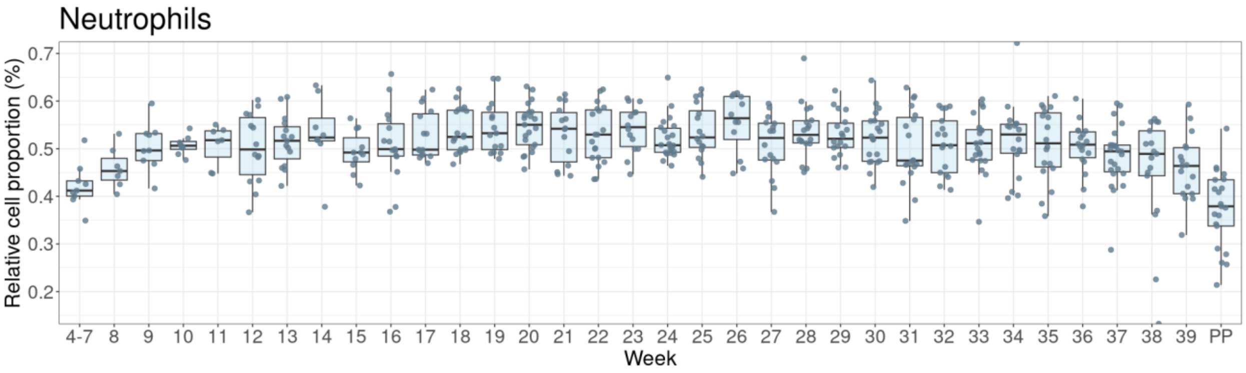
Neutrophils
Relative cell proportion (%)
Week

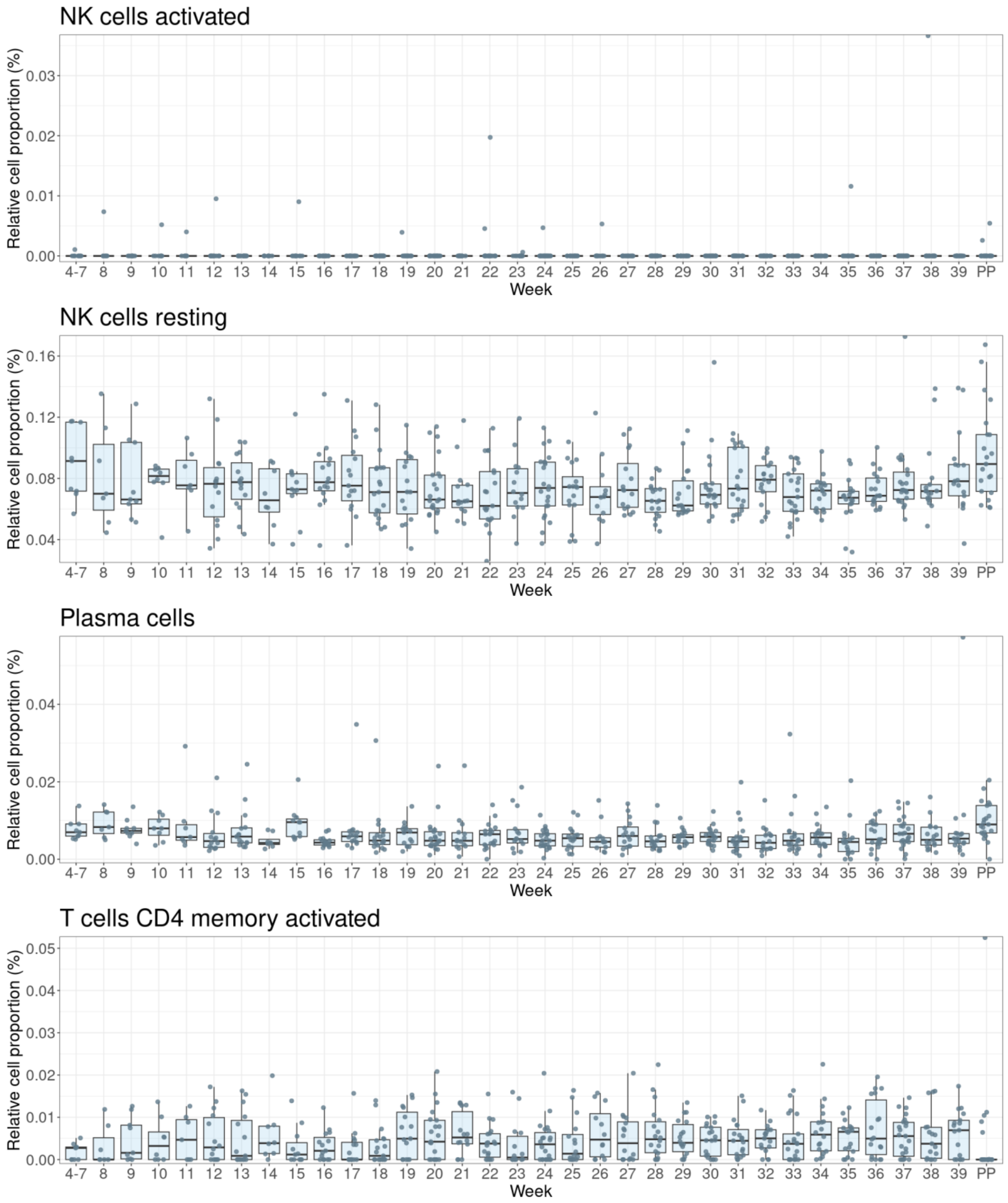
NK cells activated
Relative cell proportion (%)
0.00
0.01
0.02
0.03
4-7 8 9 10 11 12 13 14 15 16 17 18 19 20 21 22 23 24 25 26 27 28 29 30 31 32 33 34 35 36 37 38 39 PP
Week
NK cells resting
Relative cell proportion (%)
0.04
0.08
0.12
0.16
4-7 8 9 10 11 12 13 14 15 16 17 18 19 20 21 22 23 24 25 26 27 28 29 30 31 32 33 34 35 36 37 38 39 PP
Week
Plasma cells
Relative cell proportion (%)
0.00
0.02
0.04
4-7 8 9 10 11 12 13 14 15 16 17 18 19 20 21 22 23 24 25 26 27 28 29 30 31 32 33 34 35 36 37 38 39 PP
Week
T cells CD4 memory activated
Relative cell proportion (%)
0.00
0.01
0.02
0.03
0.04
0.05
4-7 8 9 10 11 12 13 14 15 16 17 18 19 20 21 22 23 24 25 26 27 28 29 30 31 32 33 34 35 36 37 38 39 PP
Week

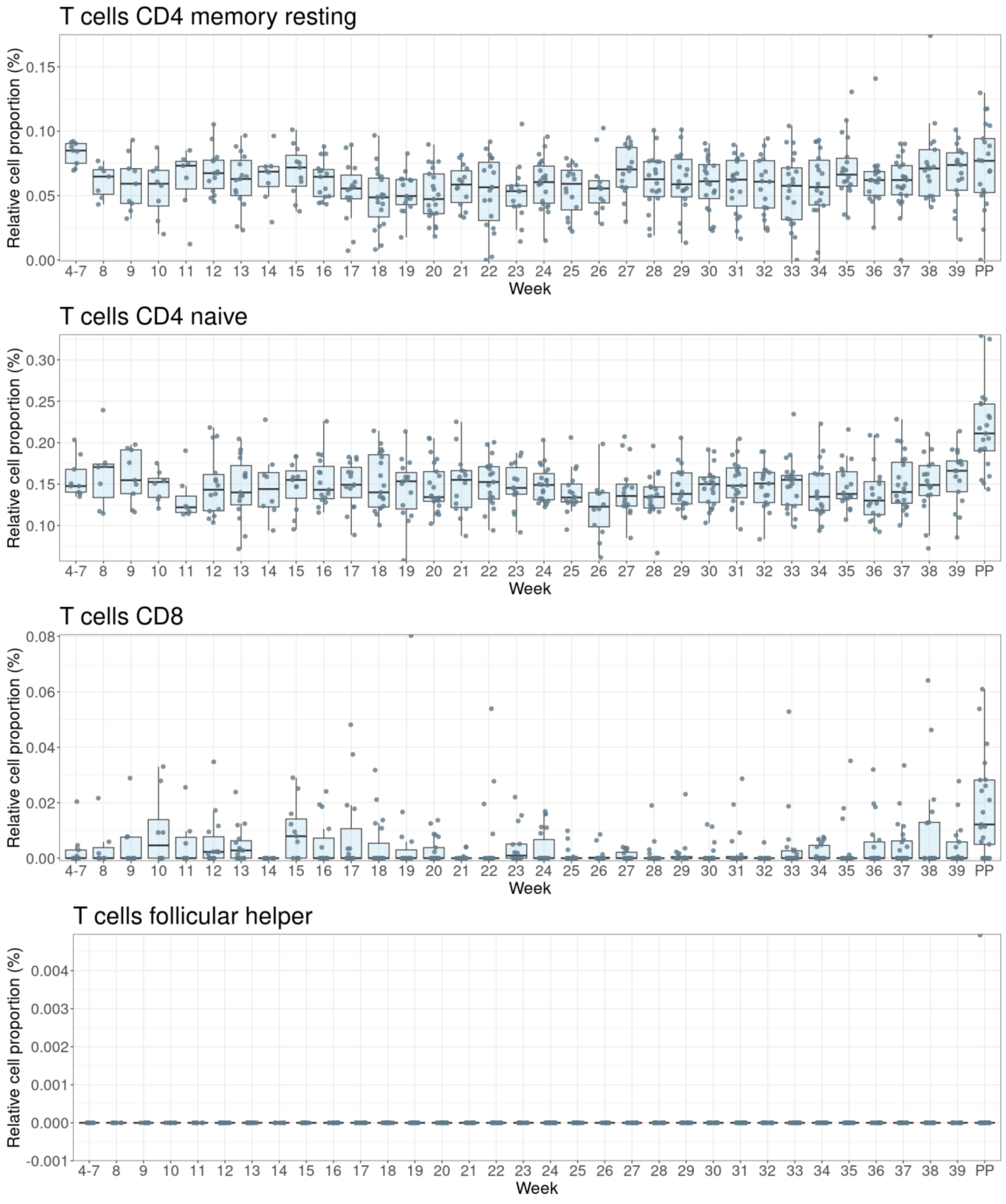

T cells CD4 memory resting
Relative cell proportion (%)
0.00
0.05
0.10
0.15
4-7 8 9 10 11 12 13 14 15 16 17 18 19 20 21 22 23 24 25 26 27 28 29 30 31 32 33 34 35 36 37 38 39 PP
Week
T cells CD4 naive
Relative cell proportion (%)
0.10
0.15
0.20
0.25
0.30
4-7 8 9 10 11 12 13 14 15 16 17 18 19 20 21 22 23 24 25 26 27 28 29 30 31 32 33 34 35 36 37 38 39 PP
Week
T cells CD8
Relative cell proportion (%)
0.00
0.02
0.04
0.06
0.08
4-7 8 9 10 11 12 13 14 15 16 17 18 19 20 21 22 23 24 25 26 27 28 29 30 31 32 33 34 35 36 37 38 39 PP
Week
T cells follicular helper
Relative cell proportion (%)
-0.001
0.000
0.001
0.002
0.003
0.004
4-7 8 9 10 11 12 13 14 15 16 17 18 19 20 21 22 23 24 25 26 27 28 29 30 31 32 33 34 35 36 37 38 39 PP
Week

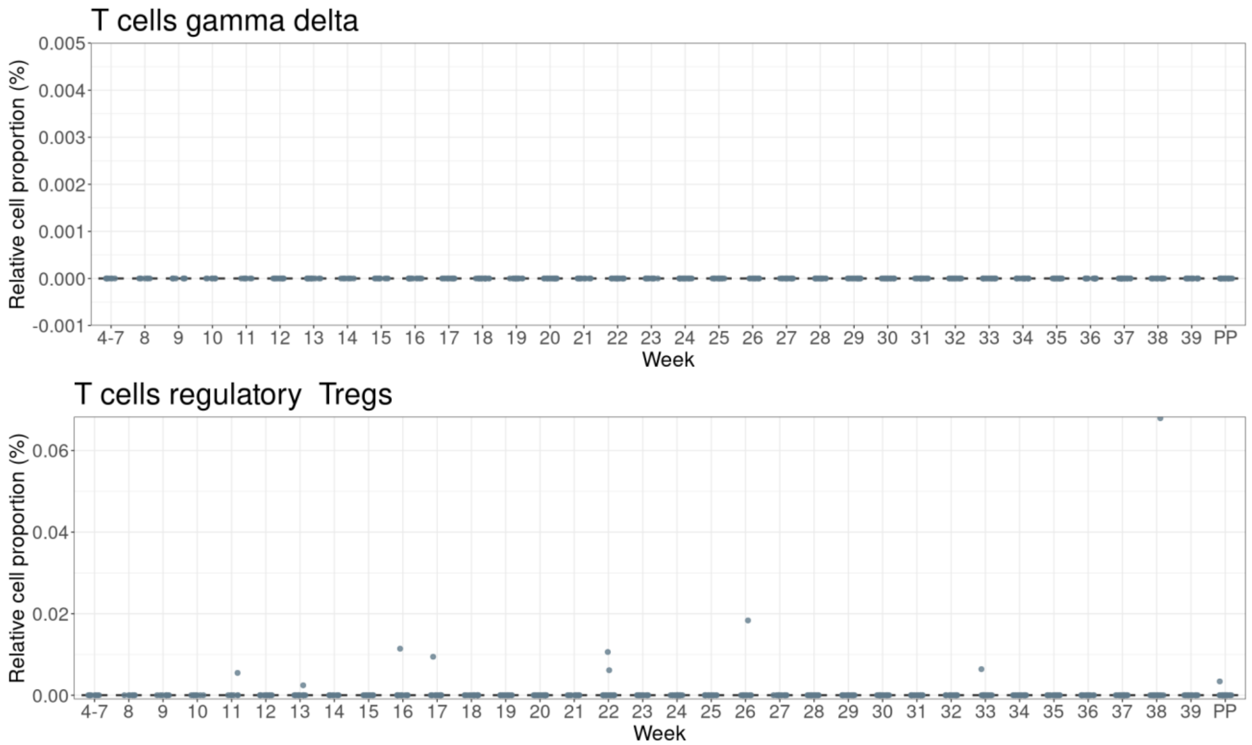

**Supplementary Fig. 3:** Box plots of the relative cell type proportions of 22 immune cells across gestational age (in weeks) obtained from the cell deconvolution analysis with CIBERSORTx. The proportions sum to one across all samples and y-axes are adjusted for each cell type.

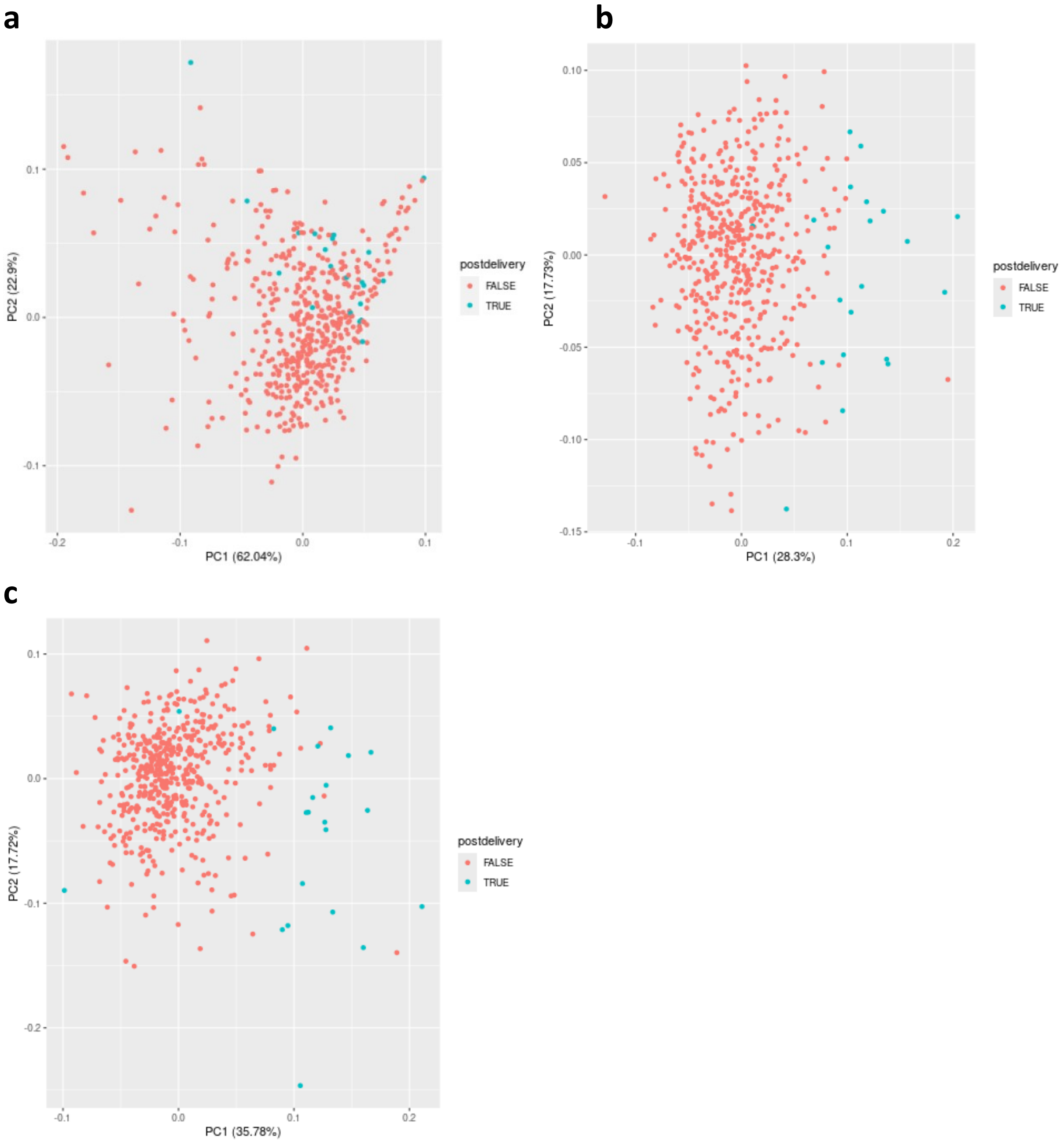

**Supplementary Fig. 4: Principal component analysis of transcript levels.** The colors indicate pregnancy vs post-partum samples**. a,** raw transcript counts. **b,** counts after VST normalization. **c,** counts after both VST and regression normalization.

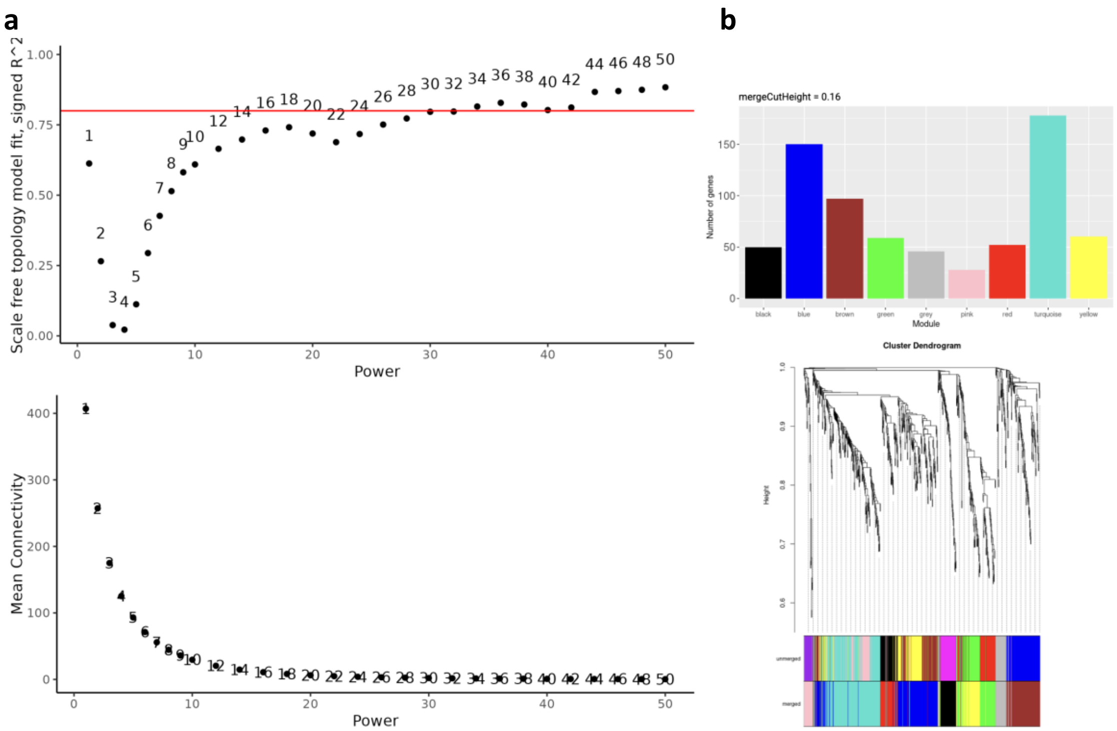

**Supplementary Fig. 5: Parameter evaluation behind the signed WGCNA analyses. a,** Scale independence (top) and mean connectivity (bottom) with selected soft power threshold 16 to maximize R^2 and minimize mean connectivity. **b,** The final module sizes (top) and cluster dendrogram (bottom) for the final network with merge cut height 0.16.

### Supplementary Tables

**Supplementary Table 1:** Results for 720 transcripts that were robustly validated in the replication and combined cohorts and the classification of the transcripts in the WGCNA derived co-expressed modules. The table contains columns corresponding to the module color, gene symbol, Ensembl id, and the model p-value of the discovery, replication and combined cohorts, respectively. The table is supplied as an online excel sheet.

**Supplementary Table 2:** Table of the transcripts found in our work, Wright et al. (2023) and Gomez-Lopez et al. (2019) with columns describing whether or not a given gene was reported in each work. Included are our p-values from the discovery, replication, and combined batches. The genes are sorted after the latter. The table is supplied as an online excel sheet.

**Supplementary Table 3:** Mapping of the 720 pregnancy associated transcripts to 664 Ensembl Protein IDs. The 56 transcripts not mapping to protein IDs primarily represented long noncoding RNAs. The table is supplied as an online excel sheet.

**Supplementary Table 4:** Tests of association between estimated cell type proportions and pregnancy week. The table shows Benjamini-Hochberg adjusted *P* values obtained from a linear mixed effects model evaluating the estimated cell type proportions of each gestational week compared to the post-partum levels. Note that the cell types T cells gamma delta and Macrophages M1 had relative proportions of zero across all gestational weeks and have thus not been modeled to obtain *P* values. The table is supplied as an online excel sheet.
